# Tumor suppressor Hypermethylated in Cancer 1 represses expression of cell cycle regulator E2F7 in human primary cells

**DOI:** 10.1101/2022.07.25.501405

**Authors:** Lucie Lanikova, Jiri Svec, Lucie Janeckova, Vendula Pospichalova, Nikol Dibus, Martina Vojtechova, Dusan Hrckulak, Eva Sloncova, Hynek Strnad, Vladimir Korinek

## Abstract

Hypermethylated in Cancer 1 (HIC1) is an established tumor suppressor, which is frequently inactivated in various cancers. In colorectal carcinoma (CRC), silencing of HIC1 has been recognized as one of the important events in malignant tumor progression. Strikingly, CRC patients with high HIC1 expression have a worse prognosis than patients with relatively low *HIC1* mRNA levels. To analyze the function of HIC1, we performed expression profiling of human primary fibroblasts after downregulation of HIC1 by RNA interference. We show that HIC1 deficiency triggers a p53-dependent response and that disruption of the *HIC1* gene in human colon cells delays cell cycle progression under serum deficiency conditions. Moreover, treatment with etoposide, a DNA-damaging agent, significantly impairs the proliferation rate and dynamics of damaged DNA repair in HIC1-deficient compared with wild-type cells. One of the genes upregulated in HIC1-depleted cells encodes cell cycle regulator E2F7. E2F7 is an atypical member of the E2F family, which functions primarily as a transcriptional repressor, and its downregulation is essential for proper cell cycle progression and expression of genes involved in DNA repair. We demonstrated that E2F7 is indeed the target of transcriptional repression mediated by HIC1. Moreover, our results suggest that the phenotypic manifestations associated with loss of the *HIC1* gene, in particular the changes in cell cycle progression and slowed repair of damaged DNA, are caused by dysregulation of E2F7 expression. Finally, we observed an inverse relationship between HIC1 and E2F7 in a panel of CRC. Importantly, CRC patients who express relatively high levels of E2F7 have a remarkably better prognosis than patients with intermediate or low levels of E2F7 expression.

## Introduction

The Hypermethylated in Cancer 1 (*HIC1*) gene is frequently lost or silenced in many human cancers, suggesting its role as a tumor suppressor (1). Functional evidence for this conclusion has been provided by gene targeting studies in mice. While mice with homozygous loss of *Hic1* exhibited multiple developmental defects leading to embryonic and perinatal lethality (2), mice with a functional *Hic1* allele were viable and fertile. However, Hic1^+/-^ animals spontaneously developed multiple tumors in various tissues later in life (3). The HIC1 protein has been characterized as a transcriptional repressor containing the N-terminal Broad complex, Tramtrack, and Bric à brac/POx viruses and zinc finger (BTB/POZ) protein-protein interacting domain responsible for HIC1 multimerization and binding with multiple partners, and the C-terminal portion containing five Krüppel-like C2H2 zinc fingers. The latter portion provides affinity for the HIC1-responsive element (HiRE) present in the regulatory regions of genes repressed by HIC1 (1). Histone deacetylase sirtuin 1 (SIRT1) was one of the proteins associated with HIC1 via the BTB/POZ domain. Interestingly, the HIC1-SIRT1 complex binds directly to the *SIRT1* promoter and represses transcription of the *SIRT1* gene. Inactivation of HIC1 attenuates SIRT1 production, leading to deacetylation of p53, followed by suppression of the proapototic response induced by DNA damage (4).

Based on the presence of HiRE in regulatory regions, several other HIC1 target genes have been identified. The genes encode various proteins involved in proliferation, differentiation and apoptosis (5-8). Another important function of HIC1 is to inhibit the transcriptional complexes that mediate STAT3 (9) and canonical Wnt signaling (10). In many clinical studies, inactivation of HIC1 correlates with a more aggressive phenotype and poorer survival in various tumor types (reviewed in (11)). Our analysis of gene expression profiles of colorectal neoplasia samples revealed that HIC1 expression was indeed decreased in precancerous stages. However, patients with tumors with relatively high *HIC1* mRNA expression in more advanced lesions, i.e., colorectal carcinomas (CRCs), had a lower survival rate than patients with CRCs with low *HIC1* expression (12).

To uncover the molecular basis of the tumor suppressive role of HIC1, we expression profiled human primary fibroblasts after *HIC1* mRNA silencing by RNA interference (RNAi). In addition, we used the clustered regularly interspaced short palindromic repeats (CRISPR)/Cas9 system to disrupt the *HIC1* gene in human colon (HC) cells. Gene set enrichment analysis (GSEA) revealed that small inhibitory (si) RNA-mediated silencing of HIC1 primarily triggers a p53-dependent transcriptional response. The cell morphology and proliferation rate of HIC1-deficient HC cells were normal, but serum starvation resulted in a decreased proliferation rate of these cells. Cell cycle analysis revealed a statistically significant accumulation of HIC1-deficient cells in G2/M phase, suggesting delayed progression through the cell cycle. In addition, treatment with etoposide, an agent that inhibits DNA topoisomerase II and leads to double-strand breaks in genomic DNA, significantly impaired the proliferation rate of HIC1 KO HC cell compared with HIC1 wild-type (WT) cells. Subsequent analysis revealed that DNA repair is impaired in the absence of HIC1.

One of the genes upregulated after HIC1 knockdown or *HIC1* gene disruption encoded transcription factor E2F7. The function of the E2F family of transcription factors is to regulate cyclin-dependent kinases (CDK) and retinoblastoma protein (RB), which control cell cycle progression. Importantly, dysregulation of E2F7 (and E2F8) leads to delayed cell cycle transition (13). We confirmed that the *E2F7* gene is indeed repressed by HIC1. Our additional results suggest that the cell cycle defects and lower DNA damage repair dynamics observed in HIC1-deficient cells are due to the dysregulation of E2F7 expression. We also analyzed the mRNA levels of *HIC1* and *E2F7* in tumor samples obtained from colorectal tumors. We observed an inverse relationship between the expression levels of *HIC1* and *E2F7* in CRC samples. Importantly, a negative correlation between the expression of these genes was observed, and, in addition, patients with high *E2F7* expression had longer survival than individuals with tumors producing increased levels of *HIC1* mRNA.

## Materials and methods

### Cell culture, CRISPR/Cas9 editing of the HIC1 locus, and chromosome analysis

Human WI38, HFF-1, and HEK293 cells were purchased from ATCC (cat. no. CCL-75, SCRL-1041, CRL-1573, respectively). Human HC cells immortalized by human telomerase reverse transcriptase (hTERT) expression were purchased from Applied Biological Materials (cat. no. T0570). The cell lines used were not further authenticated. All cell lines were regularly tested for mycoplasma using the MycoStrip™ detection system (InvivoGen, Toulouse, France); all cell lines were maintained in Dulbecco’s Modified Eagle’s Medium (DMEM) with GlutaMAX (Thermo Fisher Scientific, Waltham, MA, USA) supplemented with 10% fetal bovine serum (Thermo Fisher Scientific) and 100 U/mL penicillin and 100 μg/mL streptomycin (both from Thermo Fisher Scientific) in a humidified atmosphere containing 5% CO2 at 37°C. Knockout HIC1 HC cell lines (HIC1 KO) were generated using the lentiCRISPRv2 vector (Addgene, #52961); HIC1-specific single guide RNA (sgRNA) was designed using the CRISPR design tool available at crispr.mit.edu; see supplementary Table S1 for the list of gRNA sequences. Cells were co-transfected with the lentiCRISPRv2 vector and pARv-RFP reporter plasmid (14); the plasmids contained the corresponding DNA sequence that drives production or is recognized by the sgRNA, respectively. Red fluorescent protein (RFP)-positive cells were sorted into 96-well plates as single cells 72 hours after transfection. HIC1 knockout was validated by sequencing DNA fragments amplified from genomic DNA derived from multiple clones. Primers are listed in supplementary Table S1. For chromosome analysis, cells were fixed in suspension and maintained in Carnoy’s solution. Twenty metaphases were evaluated for each cell clone; HIC1-deficient and control HC cell clones were analyzed approximately 15 generations after *HIC1* gene/”mock” targeting. Chromosome banding was performed using the standard G-banding procedure. The chromosome number and structure were evaluated using an Olympus BX60 microscope (Olympus Czech Group, Prague, Czech Republic) and Ikaros software (MetaSystems, Altussheim, Germany). Karyotypes were assembled according to the International System for Human Cytogenomic Nomenclature (15).

### RNA interference and DNA microarray analysis

For RNAi, WI38 and HFF-1 cells were reverse transfected with Lipofectamine RNAiMax (Thermo Fisher Scientific) according to the manufacturer’s instructions with 10 nM siRNA targeting *HIC1*, HIC1 siGENOME SMART Pool (M-006532-01; Dharmacon/Horizon Discovery, Cambridge, UK) and HIC1 Pre-designed Silencer (AM16708; Ambion/Thermo Fisher Scientific)], or with scrambled control siRNA (non-targeting siRNA Pool No. 1, Dharmacon/Horizon Discovery). Total RNA was isolated using the RNeasy Plus Mini Kit (Qiagen, Germantown, MD, USA) from WI38 and HFF-1 cells harvested at 24, 48, or 72 hours after siRNA transfection. For microarray analysis, the quality of the isolated RNA was checked using Agilent Bioanalyzer 2100 (Agilent, Santa Clara, CA, USA). RNAs with an RNA integrity number (RIN) above 8 were further processed. Two biological replicates were used for each time point and siRNA treatment. RNA samples were analyzed using Human HT expression BeadChip V4 (Illumina, San Diego, CA, USA). Raw data were processed using the bead array package from Bioconductor (16). Datasets obtained with DNA microarrays were analyzed in the R environment using the Linear models for microarray data analysis (LIMMA) package (17).

### RNA isolation and reverse-transcription quantitative polymerase chain reaction (RT-qPCR)

For RT-qPCR analysis, total RNA from WI38 and HFF-1 cells was transcribed using RevertAid reverse transcriptase (Thermo Fisher Scientific); for RT-qPCR of HC HIC1 KO cell lines, RNA was isolated using TRI reagent (Sigma Aldrich/Merck; Merck Life Science, Prague, Czech Republic), and 500 ng of DNA-free RNA was reverse transcribed using the First Strand cDNA Transcriptor Synthesis Kit (Roche Life Science, Roche Diagnostics Division, Prague, Czech Republic) according to the protocol provided by the manufacturer. Frozen human samples were digested in 600 µl of lysis buffer containing green ceramic beads and the tissue was disrupted using the MagNA Lyser Instrument (Roche Life Science). Total RNA was extracted using the RNeasy Mini Kit (Qiagen) according to the manufacturer’s instructions. Complementary DNA synthesis was performed using 1 µg of total RNA, random hexamers, and RevertAid reverse transcriptase (Thermo Fisher Scientific) according to the manufacturer’s protocol. PCR reactions were performed in triplicate using the LightCycler 480 instrument (Roche Life Science). The initial denaturation was performed at 95 °C for 7 minutes (ramp rate 4.8 °C/s); the PCR reactions were run for 45 cycles with the following parameters: denaturation 95 °C/20 s (ramp rate 4.8 °C/s), annealing 61 °C/20 s (ramp rate 2.5 °C/s), extension 72 °C/20 s (ramp rate 4.8 °C/s). At the end, the reactions were cooled at 40 °C for 30 s (ramp rate 2.5 °C/s). Quantification cycle (Cq) values for each triplicate were normalized to ubiquitin B (*UBB*) mRNA levels. Reported data represent the mean of three independent experiments; T bars indicate standard deviation (SD). For statistical analysis, Student’s paired *t*-test with unequal variance was used and p-values < 0.05 were considered statistically significant. LightCycler 480 Probes Master and Universal ProbeLibrary (UPL) hydrolysis probes (Roche Life Science) were used for RT-qPCR analysis of the human samples; Cq values for each triplicate were normalized by the geometric mean of housekeeping genes *UBB* and TATA box binding protein (*TBP*). The resulting values were averaged to obtain the ΔCq values for the biological replicates. The relative mRNA abundance (ΔCq in healthy tissue - ΔCq in neoplastic tissue) was correlated with the histological grade of the tumor samples using the Spearman (ρ) and Kendall (τ) rank order coefficients. Primer pairs and UPL probe numbers are listed in supplementary Table S1.

### Proliferation assay and cell cycle analysis

The cell number and viability were determined using CellTiter-Blue reagent (Promega Corporation, Madison, WI, USA) and Perkin-Elmer Envision analyzer (PerkinElmer, Waltham, MA, USA). Results are presented as the percentage of cell growth compared to the number of cells at the time of start. For cell cycle analysis, HIC1 KO and control cells were harvested, washed with ice-cold phosphate-buffered saline (PBS), and fixed with 70% ethanol. The cell cycle was analyzed using a BD FACSCanto II flow cytometer (BD Biosciences, Becton Dickinson Czechia, Prague 4, Czech Republic) and FlowJoTM software (data were collected for at least 10,000 cells).

### Luciferase reporter assay

The assay was performed using the Dual-Glo Luciferase Assay System and a GloMax® 20/20 Luminometer (all from Promega). To test the activity of the *E2F7* promoter, a 3kb region of the promoter containing HIC1 binding sites was amplified by PCR from human genomic DNA and cloned into the pGl4.26-luc vector (Promega); primers are listed in supplementary Table S1. The HIC1 expression construct has been described previously (10). All luciferase assays were performed in triplicate and results were normalized to Renilla (pRL-TK; Promega).

### Western blotting, immunohistochemistry, and antibodies

For Western blotting, cells were harvested in RIPA buffer (Sigma-Aldrich/Merck) supplemented with a cocktail of protease inhibitors (Roche Life Science). Proteins were resolved in SDS-polyacrylamide gels and blotted onto PVDF membranes (Millipore/Merck) or nitrocellulose membranes (Bio-Rad, Prague, Czech Republic). For immunohistochemistry, 5µm sections of tissue samples fixed in formalin and embedded in paraffin were deparaffinized in xylene and rehydrated. To unmask the antigenic sites, the sections were immersed in Target Retrieval Solution (Dako/Agilent) in a steam bath. Primary rabbit polyclonal antibodies, anti-HIC1 (HPA043372, Sigma-Aldrich/Merck) and anti-E2F7 (ab56022, Abcam, Cambridge, UK), were applied to the sections at a dilution of 1:100 overnight at 4°C or 1:200 for 1 hour at room temperature. The primary antibody was detected with peroxidase-conjugated secondary anti-rabbit antibody using the EnVision + system (Dako/Agilent); the brownish color reaction was developed with 3,3’-diaminobenzidine (DAB) substrate (Vector Laboratories, Newark, CA, USA). A detailed protocol of the immunoblotting procedure has been described previously (18); antibodies anti-E2F7 (the same as above), anti-p53 (mouse monoclonal, 1C12, Cell Signaling, Danvers, MA, USA; dilution 1:500), anti-phospho-p53 (Ser 15) (rabbit polyclonal, #9284, Cell Signaling; 1:500), an anti-α-tubulin (mouse monoclonal, TU-01, EXBIO Praha, Vestec, Czech Republic; 1:1000), anti-CtBP (mouse monoclonal, E12, Santa Cruz Biotechnology, Dallas, TX, USA; 1:1000) were used. Anti β-actin (rabbit polyclonal, # A2066, Sigmal-Aldrich/Merck; 1:1000) antibody was used as an internal control. For immunofluorescence analysis of a phosphorylated variant of histone H2AX (γH2AX) and TP53-binding protein 1 (53BP1) positive foci, HIC1 KO cells were seeded on coverslips and DNA damage was induced by incubation with 10 μM etoposide (Sigma-Aldrich/Merck) for 1 hour. At indicated time intervals, cells were fixed with 4% paraformaldehyde (Electron Microscopy Sciences, Hatfield, PA, USA) and coverslips were incubated with primary (γH2AX (05-636, Millipore/Merck) and 53BP1 (MAB3802, Millipore/Merck)) and secondary (AlexaFluor-568 and AlexaFluor-488) antibodies (Thermo Fisher Scientific; a detailed protocol describing the immunofluorescence procedure can be found elsewhere (19). Images were acquired using the Olympus ScanR system equipped with 40×/1.3 UPLFN or 40×/0.9 objective (Olympus Czech Group) or Stellaris 8 Falcon with 63×/1.3 objective (4 × zoom; Leica Microsystems, Wetzlar, Germany). Nuclei were identified based on 4′,6-diamidino-2-phenylindole (DAPI; purchased from Thermo Fisher Scientific) staining before measurement of nuclear γH2AX and 53BP1 staining, and foci were identified using the Spot detection module. Image analysis was performed with at least 400 cells per given condition.

### Ethics statement, tissue samples, and public database datasets

All human sample collection procedures were performed in accordance with relevant national and EU regulations and the World Medical Association Code of Ethics (Declaration of Helsinki). The study was approved by the Ethics Committee of the Third Faculty of Medicine of Charles University in Prague (ref. no. 26/2014). Informed consent was obtained from all patients. Paired samples of normal and neoplastic colon tissue were obtained from 70 patients who had undergone either polypectomy of colon adenomas or surgical resection of sporadic CRC (details published elsewhere (12)). Datasets GSE39582 and GSE3958 were retrieved from the Gene Expression Omnibus (20). Expression data based on RNA sequencing (RNA-seq) were retrieved from the colorectal adenocarcinoma dataset (TCGA, PanCancer Atlas) available at cBioportal (21). The data were analyzed using RNA-seq by the expectation maximization (RSEM) software package (22).

### Statistical analysis and availability of expression data

Statistical tests were performed in the R environment. Expression profile data were deposited in the ArrayExpress database (https://www.ebi.ac.uk/arrayexpress/); accession number: E-MTAB-10678.

## Results

### Expression profiling of primary human cells after HIC1 silencing

We used primary human lung fibroblasts WI38 treated with HIC1-specific siRNAs to investigate changes in the expression profile caused by decreased HIC1 expression. Cells were transfected with two different *HIC1* siRNAs (labeled ‘Amb HIC1’ or ‘Dhar HIC1’) or with a non-silencing control siRNA and harvested at 24, 48 and 72 hours post-transfection. Quantitative RT-PCR analysis of cells transfected with HIC1-specific siRNAs showed a strong decrease in *HIC1* mRNA levels compared to cells transfected with the non-silencing siRNA (Fig. 1A). In addition, gradually decreasing levels of the HIC1 protein were observed at these time points (Fig. 1B; shown are blots obtained after treatment with ‘Dhar HIC1’). Subsequently, expression profiling of total RNA isolated from cells treated with HIC1-specific and control siRNAs at 48 and 72 h time points was performed. The expression levels of 167 and 144 gene probes (representing 158 and 139 annotated genes, respectively) changed significantly 48 and 72 hours after transfection with HIC1-specific siRNAs. The expression of 73 gene probes (representing 71 annotated genes) changed significantly at both time points (Fig. 1C and supplementary Tables S2 and S3).

**Figure 1.**
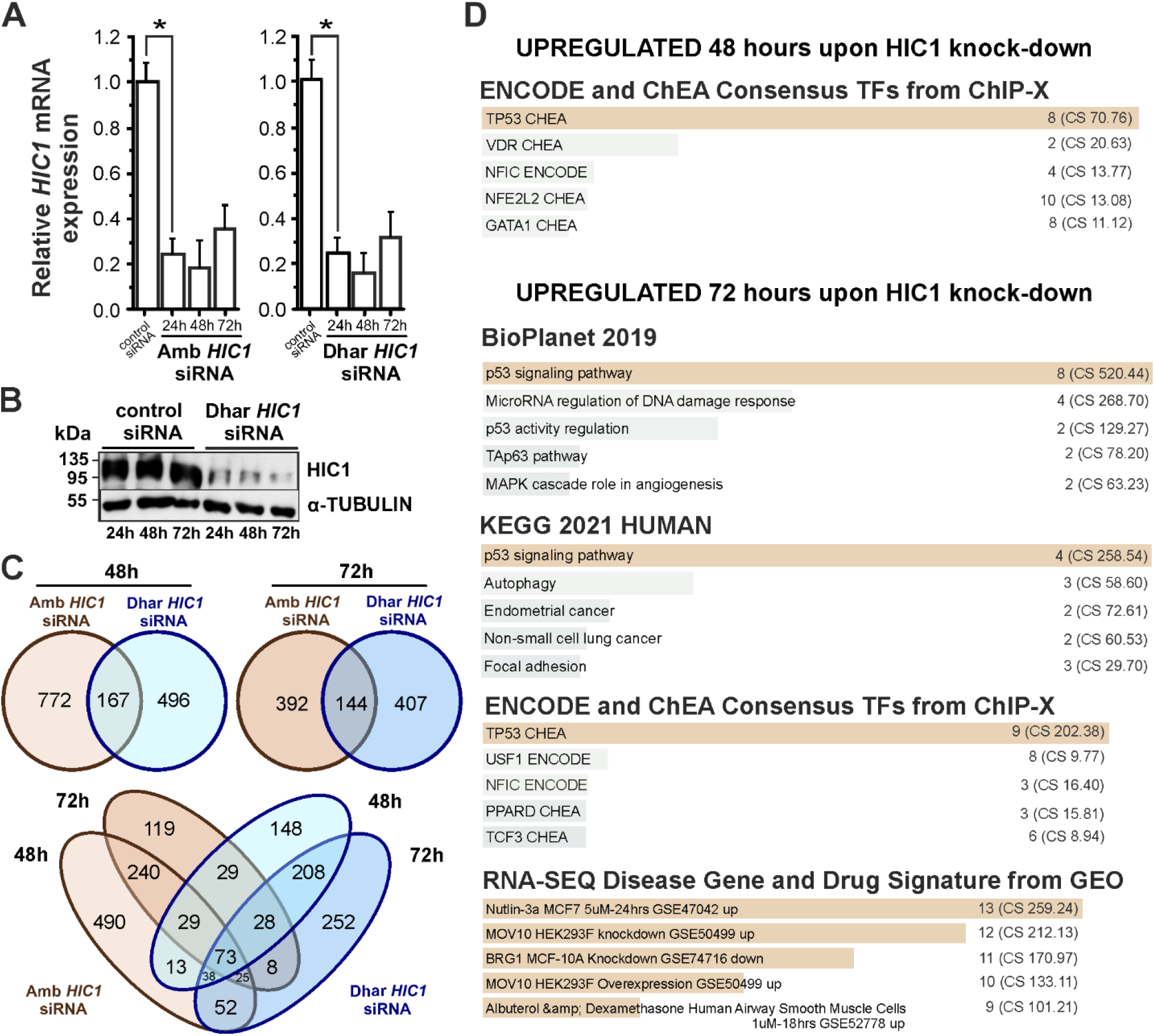
Expression profiles of human WI38 cells after knockdown of the *HIC1* gene. (A) Down-regulation of *HIC1* mRNA in WI38 cells at 24, 48 and 72 h after transfection with HIC1-specific siRNAs from Ambion (Amb HIC1 siRNA) or Dharmacon (Dhar HIC1 siRNA); transfection with non-coding siRNA was used as a control. Graphs show the results of RT-qPCR analysis of WI38 cells transfected with the indicated siRNAs; the obtained quantification cycle (Cq) values were normalized to the *UBB* mRNA levels; the *HIC1* expression level in cells treated with non-silencing control siRNA was arbitrarily set to 1. Error bars indicate standard deviations (SDs); * p-value < 0.001 (Student’s t-test). (B) Western blot analysis of HIC1 protein in WI38 cells 24, 48 and 72 hours after transfection with control or indicated HIC1-specific siRNA; immunoblotting with an anti-tubulin antibody was used as a loading control. (C) Venn diagrams showing the number of gene probes differentially expressed in WI38 cells 48 and 72 hours after transfection with HIC1-specific siRNAs compared with cells treated with control siRNA (significance criterion: q < 0.05; |log FC | ≥ 1). Annotated genes are listed in supplementary Tables S2 and S3. (D) Gene set enrichment analysis (GSEA) of genes upregulated after knockdown of HIC1; 80 and 49 genes were analyzed at the 48- and 72-hour time points, respectively, using the indicated GSEA tools. The p-value (calculated from Fisher’s exact test) was assigned to each category; five categories with the highest combined score (CS; CS was calculated by multiplying the logarithm of the p-value by the z-score of the deviation from the expected rank) for each GSEA tool are shown. The coloring of the columns corresponds to the significance of each column in the graph - the category with the adjusted p-value < 0.05 is in brown. The number of genes in each category is indicated by the number in front of the parenthesis; |log FC|, absolute value of the binary logarithm of the relative expression intensity.

Since HIC1 functions as a transcriptional repressor, we analyzed the genes whose levels increased after HIC1 silencing. We performed GSEA of 80 and 49 genes that were upregulated 48 and 72 hours after HIC1 silencing, respectively. We used BioPlanet and Kyoto Encyclopedia of Genes and Genomes (KEGG) pathway datasets included in the Enrichr web tool (23); the datasets catalog all biological pathways, their interactions and changes in various healthy or diseased conditions. The analysis revealed that the pathway most affected by the decreased expression of HIC1 was the p53 pathway (Fig. 1D). Note that the significance criterion (adjusted p-value < 0.05) for the p53-pathway dataset was reached only for the 72-hour siRNA treatment time interval. Subsequent analysis of potential binding sites of transcription factors (TFs) using the Encyclopedia of DNA Elements (ENCODE) project web tool (24) and the ChIP Enrichment Analysis (ChEA) database derived from the integration of genome-wide experiments using ChIP-Chip, ChIP-seq, ChIP-PET, and DamID (these methods were referred to as ChIP-X (25)) revealed that eight and nine genes that were upregulated 48 and 72 hours after HIC1 downregulation, respectively, were possibly regulated by p53. In addition, comparison of the same gene list with gene expression signatures from RNA-seq studies obtained after application of drugs or specific gene manipulations/disruptions (the corresponding datasets were obtained from the Gene Expression Omnibus (GEO) repository) showed the most significant match (13 genes) with the group of genes obtained after activation (stabilization) of p53 in MCF-7 breast cancer cells treated with small molecule Nutlin-3a (26) (Fig. 1D and not shown).

### E2F7 expression was suppressed by HIC1

One of the genes that was upregulated after knockdown of HIC1 in the 48- and 72-hour time interval was the gene encoding transcription factor E2F7 (supplementary Table S2). Using RT-qPCR analysis, we confirmed that the expression of *E2F7* mRNA was increased after HIC1 knockdown with both HIC1-specific siRNAs not only in WI38 cells, but also in primary human foreskin fibroblasts HFF-1 (Fig. 2A and data not shown). We then generated an E2F7-luciferase reporter (E2F7-luc) by cloning the *E2F7* promoter region containing a putative HiRE upstream of the luciferase gene and tested its response to HIC1. The reporter activity was increased after HIC1 knockout (Fig. 2B, left graph). In contrast, co-transfection of the reporter with increasing amounts of the HIC1 expression construct resulted in decreased E2F7-luc reporter activity, confirming transcriptional repression of E2F7 by HIC1 (Fig. 2B, right diagram). Furthermore, we used the CRISPR/Cas9 system to disrupt the *HIC1* gene in human HC cells (27). Alignment of the *HIC1* locus was confirmed by sequencing the corresponding genomic regions, and three HIC1 KO HC cell clones, #13, #20, and #25 (supplementary Fig. S1), were used for further analysis. The control HIC1-proficient clones transfected with the Cas9-expressing vector lacking the sgRNA expression cassette were designated CTRL #1 and CTRL #2. We encountered a problem in detecting endogenous HIC1 protein in cell lysates obtained by Western blotting from HC cells. Nevertheless, RT-qPCR analysis revealed a marked decrease in *HIC1* mRNA in HC cells transfected with HIC1-specific gRNA (Fig. 2C). Because the region spanning the translation initiation region of *HIC1* mRNA was targeted (supplementary Fig. S1), we suspected that *HIC1* mRNA levels were reduced by a nonsense-mediated RNA decay mechanism (28) (Fig. 2C).

**Figure 2.**
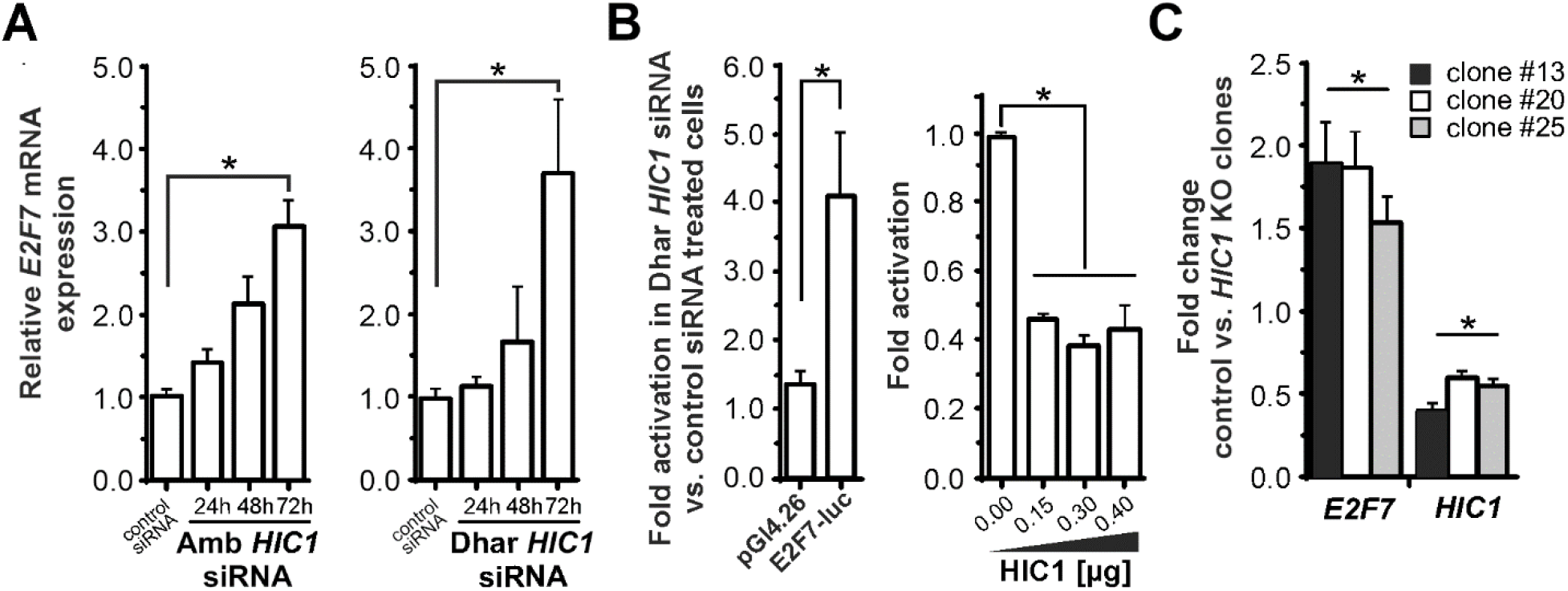
HIC1 suppresses E2F7 expression. (A) Quantitative RT-PCR of *E2F7* mRNA after HIC1 knockdown. HFF-1 cells were transfected with HIC1-specific or control non-silencing siRNA and harvested at 24, 48, and 72 hours after transfection. Results were normalized to *UBB* mRNA levels; *HIC1* expression levels in cells treated with control siRNA were set to 1. (B) Left: Luciferase reporter assay in HFF-1 cells. Cells were transfected with Dhar HIC1 or control siRNA; 48 hours later, cells were transfected with the E2F7-luc reporter or the “empty” pGl4.26 vector. Luciferase activity was determined 24 hours after the second transfection. Right: luciferase reporter assay in HEK293 cells co-transfected with the E2F7-luc reporter and increasing amounts of the HIC1-expressing vector. Luciferase reporter assays were performed in triplicate; results were normalized to the internal Renilla control. Luciferase activity in cells transfected with non-silencing siRNA (left panel) or with “empty” expression vector (panel B) was set to 1. (C) Up-regulation of E2F7 expression in HC HIC1 KO cell clones detected by RT-qPCR; results were normalized to *UBB* and *β-ACTIN* mRNA levels. The average level of *E2F7* and *HIC1* expression in two HC cell clones CTRL #1 and CTRL #2 transfected with the CRIPSR/Cas9 vector without HIC1-specific sgRNA was set to 1. Error bars indicate SDs; * p-value < 0.05 (Student’s t-test).

### E2F7 displays elevated expression in human ‘HIC-low’ CRCs

We have previously shown that the expression of HIC1 varies in CRCs (12). Therefore, we sought to analyze a possible relationship between E2F7 and HIC1 in CRC samples. To determine changes in *E2F7* expression during colorectal neoplastic progression, we analyzed samples from 70 patients (12). Interestingly, we found significant upregulation of the *E2F7* gene along the neoplastic progression stage (Fig. 3A). Moreover, RT-qPCR analysis of total RNA isolated from colorectal tumor samples showed that the increased *E2F7* expression correlated with the decline in *HIC1* mRNA (Fig. 3B). Next, we used the cBioPortal for Cancer Genomics Webtool (https://www.cbioportal.org/) to analyze the expression profile of 524 CRC samples. The RNA sequencing data showed an inverse correlation between *E2F7* and *HIC1* expression levels; this correlation was relatively weak, although significant (Fig. 3C). The inverse relationship between E2F7 and HIC1 expression was confirmed by statistical analysis of public datasets GSE37892 and GSE39582, which contain DNA microarray-based expression analyses of 130 and 443 colorectal cancer samples, respectively (29, 30) (supplementary Fig. S2). Next, we performed immunohistochemical detection of HIC1 and E2F7 in the progressive stages of colon carcinogenesis consisting of healthy mucosa, hyperplastic polyps, adenomas with low- or high-grade dysplasia, and invasive adenocarcinomas. In the healthy colonic mucosa, HIC1 expression showed an apicobasal gradient during the progression of colorectal cancer. In the healthy mucosa, nuclear and cytoplasmic E2F7 staining was observed in almost all epithelial cells with marked positivity in the superficial colonocytes; immunopositivity was maintained in the hyperplastic lesion. Starting with the low-grade adenoma, we observed heterogeneous nuclear and strong cytoplasmic E2F7 immunostaining (Fig. 3D). Finally, we investigated whether the E2F7 expression level was related to some specific clinicopathological features included in the GSE39582 dataset (30). First, we determined the relative *E2F7* mRNA expression in each CRC. Samples that fell in the first and last deciles according to E2F7 expression were defined as ‘E2F7-high’ and ‘E2F7-low’, respectively. The rest of the samples were labeled as ‘Other’; *HIC1* expression levels were reported in the same way in each CRC. Interestingly, patients with high E2F7 expression had better survival than patients with low E2F7 expression. In contrast, HIC1-high/low CRCs showed the opposite trend, in agreement with previously published results (12) (Fig. 3E).

**Figure 3.**
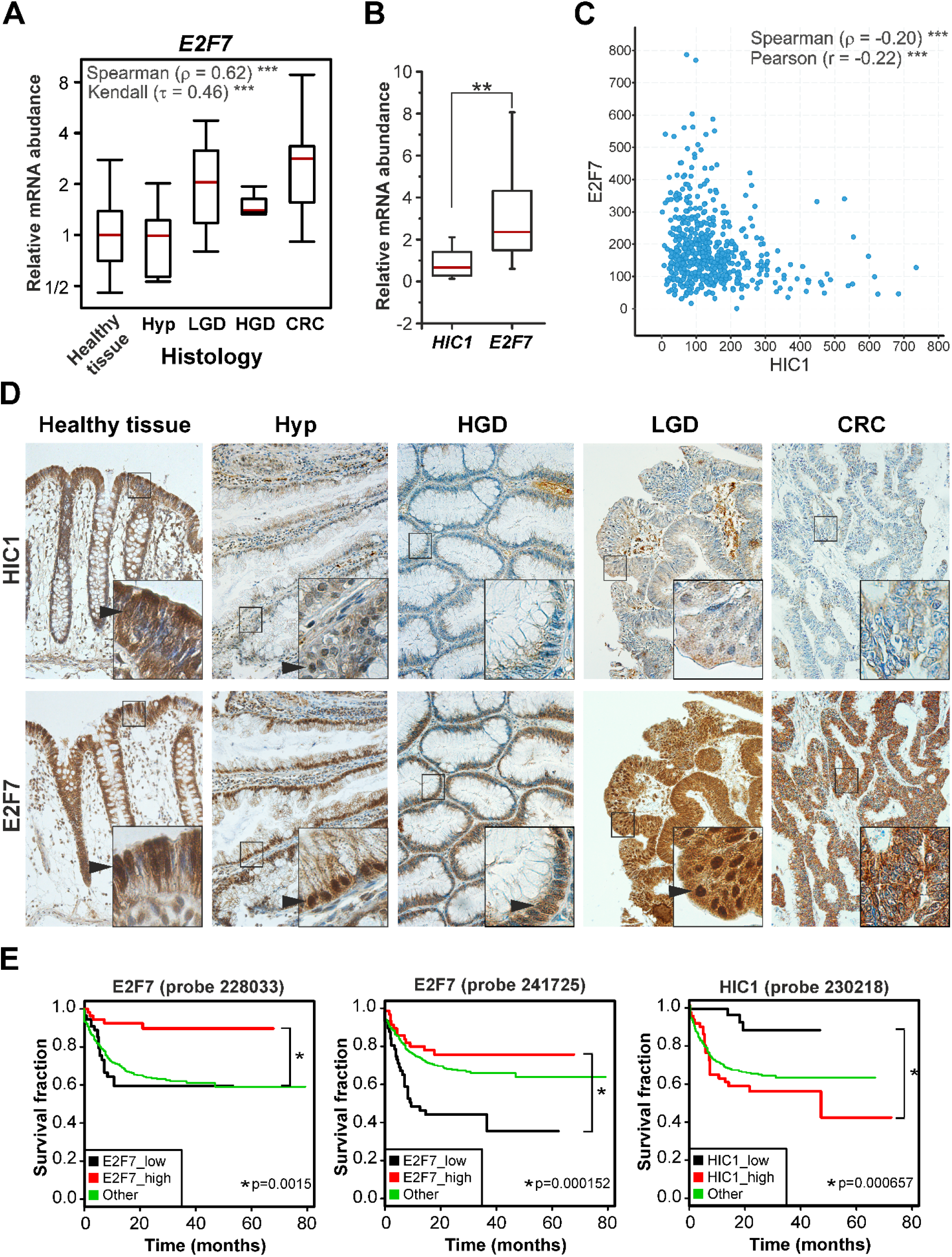
Inverse relationship between E2F7 and HIC1 expression in colorectal neoplasms. (A) *E2F7* expression changes during tumor progression. Left, RT-qPCR analysis of *E2F7* mRNA levels in healthy mucosa, hyperplastic adenomas (Hyp; n = 9), adenomas with low-grade (LGD; n = 24) or high-grade (HGD; n = 25) dysplasia, and adenoma carcinomas (CRC; n = 12). The framed areas correspond to the second and third quartiles; the median ΔCq values for each category are indicated as the red line. The association between the *E2F7* expression profile and the histology grade of the neoplasia is significant, as shown by the Spearman and Kendall coefficient values. Details of the analysis can be found in Material and methods. ***, p < 0.001; statistical significance of the correlation coefficients was estimated using the cor.test function of the R statistical environment (version 3.1.0). (B) Comparison of *HIC1* and *E2F7* expression in carcinoma samples (n = 7) by RT-qPCR. Quantitative analysis was performed as indicated in panel (A); expression levels of *HIC1* and *E2F7* mRNA in matched healthy colon mucosa were set to 1; error bars indicate SDs; *, p-value < 0.05 (Student’s t-test). (C) Inverse correlation between *E2F7* and *HIC1* expression in 524 CRC samples deposited in cBioPortal. Transcripts per million (TPM) values are indicated on the axes. The inverse relationship between the compared genes is significant, as evidenced by the Spearman and Pearson coefficient values; ***, p < 0.001. (D) Representative light microscopy images of immunohistochemical detection of HIC1 and E2F7 in healthy colon and in different histological types of colorectal neoplasms. Protein localization was visualized (brownish precipitate) using 3,3’-diaminobenzidine staining (DAB). Samples were counterstained with hematoxylin (blue nuclei); magnified images are shown in the insets. Black arrowheads indicate nuclear immunopositivity. Original magnification: 200 ×. (E) Kaplan-Meier plots of survival of patients with colorectal neoplasms based on the expression of *HIC1* and *E2F7* analyzed by RNA hybridization on DNA microarrays (corresponding gene probes are indicated); *, p < 0.05 (Student’s *t*-test).

### HIC1 knockout in HC cells disturbed cell cycle progression

Based on functional studies, E2Fs can be considered as transcriptional activators (E2F1-E2F3) or repressors (E2F4-E2F8). Production of E2F activators peaks in late G1 phase and promotes cell entry into S phase. In contrast, the level of E2F repressors reaches its maximum in S and early G2 phases (31, 32). Yuan and colleagues have shown that downregulation of E2F7 is essential for proper cell cycle progression (13). Cell morphology and proliferation of HIC1 KO cells clones were normal compared to parental HC cells or CTRL #1 and CTRL #2 cell clones (Fig. 4A, left graph, and data not shown). Therefore, we decided to expose these cells to stress conditions. Interestingly, serum deficiency-induced stress resulted in decreased cell proliferation in HIC1 KO cell clones (Fig. 4A, right diagram). Moreover, cell cycle analysis revealed statistically significant accumulation of cells in G2/M phase, indicating delayed progression through the cell cycle (Fig. 4B). This observation was rather unexpected considering the tumor suppressive role of HIC1. Recently, Szczepny and colleagues have reported that inactivation of Hic1 in MEFs leads to chromosomal instability and G2/M arrest (33). Karyotyping of HIC1 KO and control HC cell clones revealed an increased number of aberrant karyotypes in HIC1-deficient cells. However, chromosomal aberrations, albeit to a lesser extent, were also observed in control cells (supplementary Fig. S3). Thus, it cannot be said with certainty whether the observed chromosomal instability is due to the loss of *HIC1* gene or to an intrinsic property of HC cells. Subsequently, we treated HC cells with etoposide, which inhibits DNA topoisomerase II, leading to double-strand breaks in replicating genomic DNA (34). The proliferation rate of HIC1 KO cell clones was severely impaired in the presence of the agent in comparison to the control clones with the intact *HIC* gene (Fig. 5A). Next, we stained γH2AX- and 53BP1-positive foci formation in control and HIC1 KO cell clones after etoposide treatment (Fig. 5B; fluorescent microphotographs stained for γH2AX are shown). Subsequent quantification indicated increased numbers of the nuclear foci in HIC1 KO cells not only at different time points, but also in untreated HIC1-deficient cells, suggesting impaired DNA repair in the HIC1 absence (Figure 2C; data for HIC1 KO clone #20 and control clone #1 are shown). Finally, we treated HC cells with increasing concentrations of etoposide and quantified the E2F7 expression levels. As shown in Figure 5D, in HIC1-proficient HC cells, *E2F7* mRNA was decreased upon etoposide treatment (when compared to untreated cells). However, in HIC1 KO cell clones, E2F7 transcription was increased.

**Figure 4.**
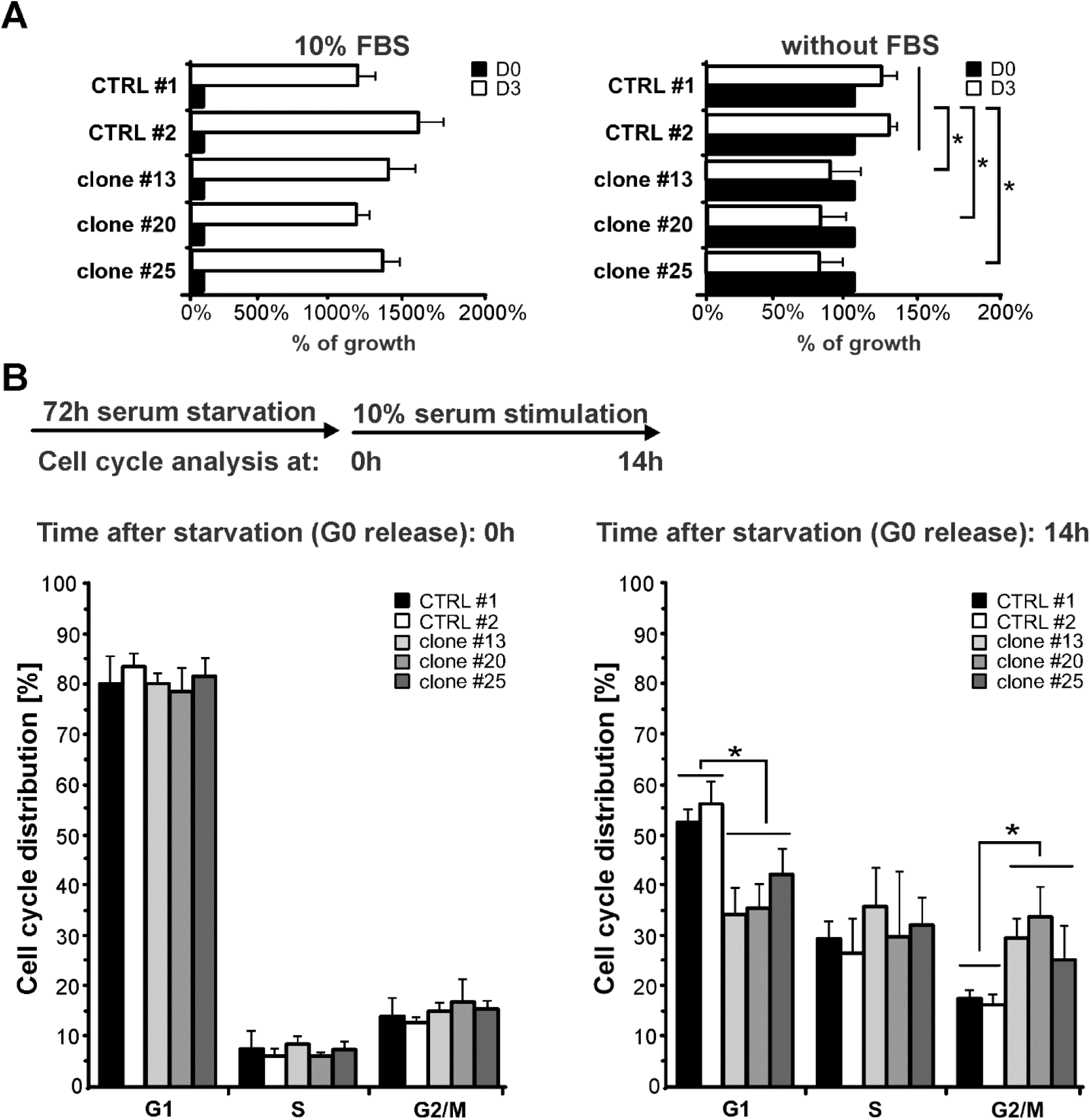
Knockout of HIC1 in HC cells disrupts cycle progression under stress conditions. (A) Proliferation rates of HIC1 KO HC cell clones #13, #20 and #25 and “mock-edited” HC cells (CTR L#1 and CTRL #2) under normal culture conditions (left) and under serum starvation for 72 hours (right). Results are presented as the percentage of cell growth at the 72-hour time point (D3); the number of cells at the time of serum starvation (D0) was set to 100%. (B) Cell cycle distribution of HIC1-proficient and HIC1 KO HC cells after 72-hour serum withdrawal (left) and 14 hours after release to the cell cycle (right). The top schematic shows the timing of the experiment; error bars indicate SD; * p-value < 0.05 (Student’s t-test).

**Figure 5.**
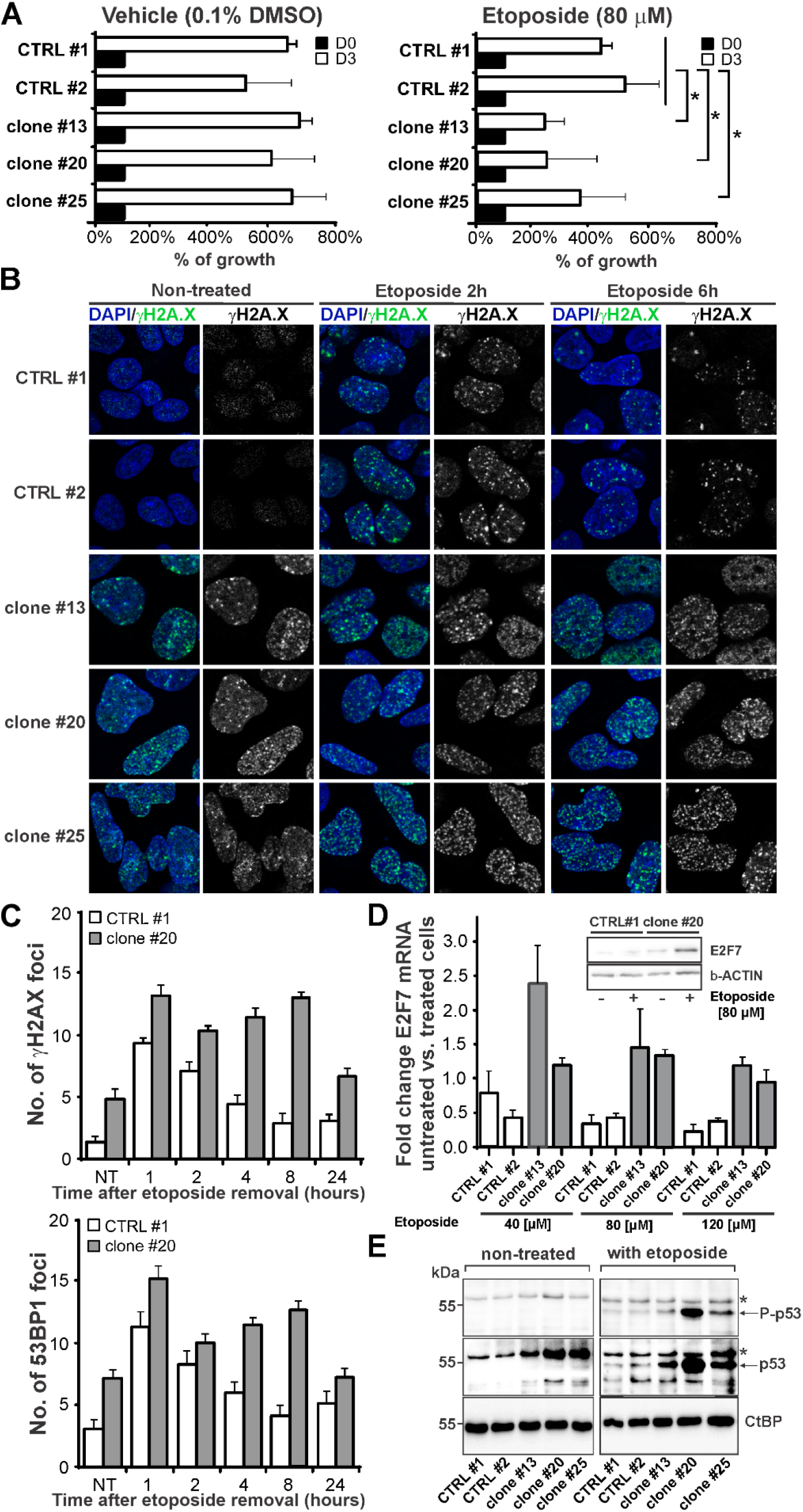
HIC1 knockout affects recovery from DNA damage. (A) Proliferation rate of HIC1 KO HC cell clone #13, #20, and #25) and control clones (CTRL #1 and #2) after etoposide (80 µ M) treatment for 72 hours. The results are demonstrated as a percentage of cell growth in comparison to the number of cells at time D0 (set to 100%). Error bars indicate SD; * p-value < 0.05 (Student’s t-test). (B) Representative fluorescent images of the γH2AX-positive nuclear foci present in control and HIC1-deficient HC cells. Cells were treated with etoposide (10 μM) for 1 hour (or left untreated) and then analyzed at the indicated time points after etoposide removal. (C) Quantification of γH2AX (upper diagram) and 53BP1 (lower diagram) foci in clone #20 and CTRL #1 HC cells after etoposide (10 µ M) treatment for 1 hour. Cells were fixed after pre-extraction at indicated time points and stained with a γH2AX and 53BP1 antibody. The mean of median foci number +/-SD is plotted (n ≥ 3). (D) Upregulation of *E2F7* mRNA level in HIC1 KO HC cells after treatment with etoposide. Indicated control and HIC1 KO cell clones were treated with increasing etoposide concentrations for 24 hours, harvested and analyzed by RT-qPCR or Western blotting (inset). The results of RT-qPCR were normalized to *UBB* and *β-ACTIN* mRNA levels. *E2F7* expression in cells treated with the solvent (DMSO) only were set to 1; immunoblotting with an anti-β-ACTIN antibody was used as a loading control. (E) Western blot analysis of p53 protein in HIC1-deficient and control HC cells. Cells were treated with etoposide (80 µ M) for 24 hours or left untreated. Whole-cell lysates were immunoblotted with an antibody that recognizes the entire pool of p53 protein (p53) or its transcriptionally active form phosphorylated on serine 15 (P-p53); the asterisk indicates a nonspecific band; CtBP, loading control; 53BP1, TP53-binding protein 1; γH2AX, a phosphorylated variant of histone H2AX.

The expression profile in human primary cells after suppression of *HIC1* gene expression was dominated by genes identified as target genes of the p53 protein. At the same time, Hic1-deficient HC cells produced markers of damage even without incubation with etoposide. Therefore, we subsequently tested p53 protein levels in HIC1-deficient and control HC cells; immunoblotting was also performed with an antibody that recognizes transcriptionally active p53 protein phosphorylated at serine 15 (35). As shown in Fig. 5E, we did not observe stabilization of p53 protein in cells that had not been incubated with etoposide. In addition, HIC1-deficient HC cells exhibited increased levels of p53 protein as well as its form phosphorylated at serine 15 after 24 hours of exposure to etoposide.

## Discussion

Hypermethylated in cancer 1 is an established tumor suppressor, which is frequently inactivated in various cancers including CRC. In the conventional model, HIC1 functions as a transcriptional repressor. The loss of HIC1 by gene silencing or chromosomal rearrangement results in aberrant expression of cancer-associated (driver) genes. To identify genes that are repressed by HIC1, we transiently downregulated *HIC1* mRNA by RNAi. For the actual analysis of the effects of *HIC1* gene loss in the context of intestinal epithelial cells, we used immortalized cells derived from the human colon in which the *HIC1* gene was disrupted by the CRISPR/Cas9 system.

Surprisingly, we found no significant overlap between our gene sets and the gene signatures obtained by expression profiling of MEFs after deletion of the *Hic1* allele, as previously published in two independent studies (33, 36). This fact likely indicates the cell specificity of HIC1-regulated genes and/or the manner in which HIC1 expression was repressed, i.e., gene inactivation of the conditional *Hic1* allele in mouse cells compared to siRNA-mediated knockdown in human cells. In addition, the *HIC1* gene did not meet the significance criteria, although RT-qPCR analysis of total RNA prior to microarray-based gene expression profiling clearly showed robust downregulation of *HIC1* mRNA. We suspected that the observed discrepancy might be caused by the very high GC content of *HIC1* mRNA leading to the low signal on the DNA chip (37).

The loss of the *HIC1* gene had a minimal effect on the cell “behavior” of HC cells. However, growth in serum-free medium resulted in lower proliferation of these cells. Subsequent cell cycle analysis showed that cells lacking HIC1 have tendency to accumulate in the G2/M phase. This observation was rather unexpected considering the tumor suppressive role of HIC1. On the other hand, Kumar described a similar effect of HIC1 depletion in glioma cells (38). In addition, we treated HC cells with etoposide, which induces double-strand breaks in replicating cells that trigger activation of ataxia telangiectasia mutated (ATM) kinase, leading to phosphorylation of multiple proteins such as H2AX and 53BP1 (39). Culture with etoposide caused a decreased proliferation rate in HIC1-deficient cells, which was probably related to lower dynamics of repair of damaged DNA. This effect of HIC1 loss is consistent with the observation of Dehennaut and colleagues, who found that HIC1 is involved in DNA repair (8). Szczepny and colleagues recently reported that inactivation of Hic1 in MEFs leads to chromosomal instability and G2/M arrest (33). In relation to the above observation, we performed karyotype analysis of HC cells. HIC1-deficient cells indeed exhibit increased chromosomal aberrations. However, a clone of control HC cells was also chromosomally unstable. Therefore, it is not possible to say whether the chromosomal aberrations observed in HIC1-deficient HC cells are indeed directly related to HIC1 loss. Another remarkable effect of the *HIC1* gene loss was an apparent activation of the DNA damage response (DDR) even in cells cultured under standard conditions, i.e., without etoposide.

One of the genes whose expression was increased in WI38 and HFF-1 cells treated with HIC1-specific siRNAs and in HIC1-deficient HC cells was *E2F7*. Transcription factor E2F7 is an atypical member of the E2F family. E2F7 acts primarily as a transcriptional repressor that antagonizes the action of the “classical” E2F proteins, e.g., E2F1 and E2F2 (31). Interestingly, the expression of genes involved in DDR and DNA repair is regulated during the cell cycle and shows the highest expression at the G1/S transition and then decreases. The decrease in expression of these genes then depends on the transcriptional repressive function of E2F7 (40). Another interesting fact is that E2F7 levels are regulated at the posttranslational level during the cell cycle by ubiquitination and subsequent degradation after interaction with cyclin F, a substrate-recognizing component of the S phase kinase-associated protein 1 (SKP1)–cullin 1 (CUL1)–F-box protein complex (SCF). SCF is a multiprotein E3 ubiquitin ligase that controls the transition between G1/S and G2/M phases and regulates the cell cycle by targeting a number of key cell cycle regulators for proteasomal degradation (41). Knockdown of cyclin F or the E2F7 mutant, which cannot interact with cyclin F and therefore remains stable during G2 phase, results in a delay of the G2/M transition accompanied by decreased expression of genes involved in DNA repair (42). Based on our results, we conclude that the cell cycle perturbations and lower dynamics of damaged DNA repair observed in HIC1-deficient cells are possibly caused by dysregulated E2F7 expression. More specifically, E2F7 accumulation interferes with G2/M progression and/or affects the expression of genes involved in recovery after DNA damage.

Why do p53-dependent genes dominate the expression signature of HIC1 siRNA-treated primary cells? Especially considering that the loss of HIC1 increases the level of SIRT1, which deacetylates (and inactivates) p53 (4). However, some recent results suggest (in agreement with our observation) that inhibition of HIC1 leads to an increase in total p53 levels, possibly accompanied by activation of the p53-dependent response (8, 38). It should be noted that analysis of the transcriptional regulatory function of HIC1 is complicated by the fact that the genes regulated by HIC1 and p53 (partially) overlap (8). Therefore, without detailed analysis, it is not possible to confirm that a “gene of interest” is indeed the target of HIC1. It is also evident that p53 in the ‘first line’ blocks the transformation of HIC1-deficient cells. It should be emphasized that the human *HIC1* and *TP53* genes (the latter gene encodes p53) are located in the same chromosomal region. Since the region is rearranged or lost in many human cancers, the condition of loss of both genes is often met. In addition, the exact nature of the cellular stress that triggers the p53-dependent response in human cells with suppressed *HIC1* gene expression remains to be determined. Finally, the HIC1 expression is upregulated by E2F1 (43), indicating a more complex interplay between HIC1 and E2F cell cycle regulators. Our data suggest that exploitation of the HIC1-E2F7 relationship may influence the sensitivity of cancer cells to treatment.

## Abbreviations

Cq: quantification cycle
CRISPR: clustered regularly interspaced short palindromic repeats
CRC: colorectal carcinoma
GSEA: gene set enrichment analysis
HC cells: human colon cells
sgRNA: single guide RNA
HIC1: hypermethylated in cancer 1
HiRE: HIC1 responsive element
RNAi: RNA interference
RNA-seq: RNA sequencing
SD: standard deviation
siRNA: small inhibitory RNA
TFs: transcription factors
UPL: Universal ProbeLibrary

## Availability of data and materials

All data generated or analyzed during this study are included in this published article (and in the supplementary files). Raw expression profile data were deposited in the ArrayExpress database (https://www.ebi.ac.uk/arrayexpress/); accession number: E-MTAB-10678.

## Authors’ contributions

LL and VK designed the outline of the study. All authors (LL, JS, LJ, VP, NB, MV, DH, ES, HS, and VK) conducted experiments and data analysis. LL, JS, LJ, VP, and VK were involved in the preparation of the manuscript. LL and VK wrote the draft of the manuscript. VK supervised the study and wrote the final version of the manuscript. All authors have read and approved the final manuscript.

## Ethics approval and consent to participate

The study was conducted in accordance with the Declaration of Helsinki. The study was approved by the Ethics Committee of the Third Faculty of Medicine of Charles University in Prague (ref. 26/2014). Informed consent was obtained from all subjects involved in the study.

## Patient consent for publication

Not applicable.

## Competing interests

All authors declare that they have no competing financial or non-financial interests that may have influenced the conduct or presentation of the work described in this manuscript.

## Acknowledgments

We thank Michal Kolar for help with biostatistical analysis, Katerina Vadinska for histochemistry of healthy tissue and tumor samples, and Sarka Takacova for critical reading of the manuscript.

## Funding

This research was funded by the Czech Science Foundation, grant no. 18-26324S, the project National Institute for Cancer Research (Programme EXCELES, ID Project No. LX22NPO5102 - Funded by the European Union - Next Generation EU), and institutional grant no. RVO 68378050. HS was supported by the project “Center for Tumor Ecology - Research of the Cancer Microenvironment Supporting Cancer Growth and Spread” (grant no. CZ.02.1.01/0.0/0.0/16_019/0000785) financed from the Operational Program “Research, Development and Education.”

## Notes

### Competing Interest Statement

The authors have declared no competing interest.

## References

1. Wales M, Biel M, el Deiry W, et al.: p53 activates expression of HIC-1, a new candidate tumour suppressor gene on 17p13.3. Nat. Med. 1: 570–577, 1995.

2. Carter M, Johns M, Zeng X, et al.: Mice deficient in the candidate tumor suppressor gene Hic1 exhibit developmental defects of structures affected in the Miller-Dieker syndrome. Hum Mol Genet 9: 413–419, 2000.

3. Chen W, Zeng X, Carter M, et al.: Heterozygous disruption of Hic1 predisposes mice to a gender-dependent spectrum of malignant tumors. Nat. Genet. 33: 197–202, 2003.

4. Chen W, Wang D, Yen R, Luo J, Gu W and Baylin S: Tumor suppressor HIC1 directly regulates SIRT1 to modulate p53-dependent DNA-damage responses. Cell 123: 437–448, 2005.

5. Briones V, Chen S, Riegel A and Lechleider R: Mechanism of fibroblast growth factor-binding protein 1 repression by TGF-β. Biochem. Biophys. Res. Commun. 345: 595–601, 2006.

6. Vilgelm A, Hong S-M, Washington M, et al.: Characterization of ΔNp73 expression and regulation in gastric and esophageal tumors. Oncogene 29: 5861–5868, 2010.

7. Van Rechem C, Boulay G, Pinte S, Stankovic-Valentin N, Guérardel C and Leprince D: Differential Regulation of HIC1 Target Genes by CtBP and NuRD, via an Acetylation/SUMOylation Switch, in Quiescent versus Proliferating Cells. Mol Cell Biol 30: 4045–4059, 2010.

8. Dehennaut V, Loison I, Dubuissez M, Nassour J, Abbadie C and Leprince D: DNA double-strand breaks lead to activation of Hypermethylated in Cancer 1 (HIC1) by SUMOylation to regulate DNA repair. J. Biol. Chem. 288: 10254–10264, 2013.

9. Lin Y-M, Wang C-M, Jeng J-C, Leprince D and Shih H-M: HIC1 interacts with and modulates the activity of STAT3. Cell cycle (Georgetown, Tex.) 12: 2266–2276, 2013.

10. Valenta T, Lukas J, Doubravska L, Fafilek B and Korinek V: HIC1 attenuates Wnt signaling by recruitment of TCF-4 and β-catenin to the nuclear bodies. The EMBO journal 25: 2326–2337, 2006.

11. Rood B and Leprince D: Deciphering HIC1 control pathways to reveal new avenues in cancer therapeutics. Expert Opin. Ther. Targets 17: 811–827, 2013.

12. Janeckova L, Kolar M, Svec J, et al.: HIC1 expression distinguishes intestinal carcinomas sensitive to chemotherapy. Transl. Oncol. 9: 99–107, 2016.

13. Yuan R, Liu Q, Segeren H, et al.: Cyclin F-dependent degradation of E2F7 is critical for DNA repair and G2-phase progression. The EMBO Journal 38: e101430, 2019.

14. Kasparek P, Krausova M, Haneckova R, et al.: Efficient gene targeting of the Rosa26 locus in mouse zygotes using TALE nucleases. FEBS Lett 588: 3982–3988, 2014.

15. McGowan-Jordan J, Simons A and Schmid M: ISCN 2016: an international system for human cytogenomic nomenclature Reprint of: Cytogenetic and Genome Research 149, 2016.

16. Storey JD and Tibshirani R: Statistical methods for identifying differentially expressed genes in DNA microarrays. Methods Mol Biol 224: 149–157, 2003.

17. Kerr MK: Linear models for microarray data analysis: hidden similarities and differences. J Comput Biol 10: 891–901, 2003.

18. Lukas J, Mazna P, Valenta T, et al.: Dazap2 modulates transcription driven by the Wnt effector TCF-4. Nucleic Acids Res 37: 3007–3020, 2009.

19. Burdova K, Storchova R, Palek M and Macurek L: WIP1 Promotes Homologous Recombination and Modulates Sensitivity to PARP Inhibitors. Cells 8: 1258, 2019.

20. Barrett T, Troup D, Wilhite S, et al.: NCBI GEO: Archive for functional genomics data sets - Update. Nucleic acids research 39: D1005–1010, 2011.

21. Gao J, Aksoy B, Dogrusoz U, et al.: Integrative Analysis of Complex Cancer Genomics and Clinical Profiles Using the cBioPortal. Science signaling 6: pl1, 2013.

22. Li B and Dewey CN: RSEM: accurate transcript quantification from RNA-Seq data with or without a reference genome. BMC Bioinformatics 12: 323, 2011.

23. Chen E, Tan C, Kou Y, et al.: Enrichr: interactive and collaborative HTML5 gene list enrichment analysis tool. BMC Bioinform. 14: 128, 2013.

24. Davis C, Hitz B, Sloan C, et al.: The Encyclopedia of DNA elements (ENCODE): data portal update. Nucleic acids research 46: D794–D801, 2017.

25. Lachmann A, Xu H, Krishnan J, Berger S, Mazloom A and Ma’ayan A: ChEA: Transcription Factor Regulation Inferred from Integrating Genome-Wide ChIP-X Experiments. Bioinformatics (Oxford, England) 26: 2438–2444, 2010.

26. Janky R, Verfaillie A, Imrychova H and B VdS: iRegulon: from a gene list to a gene regulatory network using large motif and track collections PloS Comput Biol 10: e1003731, 2014.

27. Burocziova M, Burdova K, Martinikova AS, et al.: Truncated PPM1D impairs stem cell response to genotoxic stress and promotes growth of APC-deficient tumors in the mouse colon. Cell Death Dis 10: 818, 2019.

28. Nickless A, Bailis JM and You Z: Control of gene expression through the nonsense-mediated RNA decay pathway. Cell Biosci 7: 26, 2017.

29. Laibe S, Lagarde A, Ferrari A, Monges G, Birnbaum D and Project S: A Seven-Gene Signature Aggregates a Subgroup of Stage II Colon Cancers with Stage III. Omics : a journal of integrative biology 16: 560–565, 2012.

30. Marisa L, Reyniès Ad, Duval A, et al.: Gene Expression Classification of Colon Cancer into Molecular Subtypes: Characterization, Validation, and Prognostic Value. 10: e1001453, 2013.

31. Kent L and Leone G: The broken cycle: E2F dysfunction in cancer. Nat Rev Cancer 19: 326–338, 2019.

32. Chen H, Tsai S and Leone G: Emerging roles of E2Fs in cancer: an exit from cell cycle control. Nat Rev Cancer 9: 785–797, 2009.

33. Szczepny A, Carey K, McKenzie L, et al.: The tumor suppressor Hic1 maintains chromosomal stability independent of Tp53. Oncogene 37: 1939–1948, 2018.

34. Long B, Musial S and Brattain M: Single- and double-strand DNA breakage and repair in human lung adenocarcinoma cells exposed to etoposide and teniposide. Cancer Research 45: 3106–3112, 1985.

35. Loughery J, Cox M, Smith LM and Meek DW: Critical role for p53-serine 15 phosphorylation in stimulating transactivation at p53-responsive promoters. Nucleic Acids Res 42: 7666–7680, 2014.

36. Janeckova L, Pospichalova V, Fafilek B, et al.: HIC1 Tumor Suppressor Loss Potentiates TLR2/NF-kappaB Signaling and Promotes Tissue Damage-associated Tumorigenesis. Mol Cancer Res 13: 1139–1148, 2015.

37. Veal CD, Freeman PJ, Jacobs K, et al.: A mechanistic basis for amplification differences between samples and between genome regions. BMC Genomics 13: 455, 2012.

38. Kumar S: P53 induction accompanying G2/M arrest upon knockdown of tumor suppressor HIC1 in U87MG glioma cells. Molecular and cellular biochemistry 395: 281–290, 2014.

39. Montecucco A, Zanetta F and Biamonti G: Molecular mechanisms of etoposide. EXCLI J 14: 95–108, 2015.

40. Mitxelena J, Apraiz A, Vallejo-Rodriguez J, et al.: An E2F7-dependent transcriptional program modulates DNA damage repair and genomic stability. Nucleic Acids Res 46: 4546–4559, 2018.

41. Senft D, Qi J and Ronai ZA: Ubiquitin ligases in oncogenic transformation and cancer therapy. Nat Rev Cancer 18: 69–88, 2018.

42. Yuan R, Liu Q, Segeren HA, et al.: Cyclin F-dependent degradation of E2F7 is critical for DNA repair and G2-phase progression. EMBO J 38: e101430, 2019.

43. Jenal M, Trinh E, Britschgi C, et al.: The Tumor Suppressor Gene Hypermethylated in Cancer 1 Is Transcriptionally Regulated by E2F1. Mol Cancer Res 7: 916–922, 2009.

