## Supplementary Figures and Tables for "Tumor suppressor Hypermethylated in Cancer 1 represses expression of cell cycle regulator E2F7 in human primary cells"

#### Supplementary Figures and Tables Legends

**Figure S1.** Sequences of the CRISPR/Cas9-tergeted region and allelic changes found in the *HIC1*-locus targeted clones. Numbers indicate the positions in the *HIC1* cDNA deposited in the GenBank database (accession No. NM_006497); the CRISPR guide sequence is in bold, the translation initiation codon is underlined.

**Figure S2.** Inverse correlation of E2F7 and HIC1 expression in colorectal cancer. The data were retrieved from publicly available DNA microarray-based analyses of gene expression profiling of 196 stage II and III colon carcinomas (the upper data set) (1) and 416 colon cancers (lower dataset) (2). Both dataset included two HIC1 (Nos.: 208461_at and 230218_at) and two E2F7 probes (Nos.: 228033_at and 241725_at). Expression values are expressed as $\log_2$ of the normalized intensity of the signal from the labeled RNA hybridized to the corresponding gene probe.

**Figure S3.** Increased number of aberrant karyotypes in HIC1-deficient HC cells. For each cell clone indicated, 20 metaphases were evaluated; the graph indicates the presence of a "marker chromosome", i.e., a rearranged chromosome whose genetic origin (based on its G-banded morphology) is unknown.

**Table S1.** List of primers and probes used in the study

**Table S2.** Genes differentially expressed in WI38 cells treated with HIC1 siRNA 48 or 72 hours after transfection. Indicated are 158 and 139 genes differentially expressed in cells treated with HIC1-specific siRNA compared with control siRNA 48 or 72 hours after transfection, respectively (selection criterion: adjusted p value $< 0.05$ and $|\text{Log FC}| \geq 1$).

**Table S3. Genes differentially expressed in WI38 cells treated with HIC1 siRNA at 48- and 72-hour time intervals.** A combined list of 71 genes differentially expressed in WI38 cells treated with HIC1-specific siRNA from Ambion and Dharmacon companies vs. control siRNA at both time intervals after transfection; selection criterion: adjusted p-value $< 0.05$, $|\text{Log FC}| >= 1$.

##### Supplementary References

1. Laibe S, Lagarde A, Ferrari A, et al.: A seven-gene signature aggregates a subgroup of stage II colon cancers with stage III. OMICS 16: 560-565, 2012.
2. Marisa L, Reyniès Ad, Duval A, et al.: Gene Expression Classification of Colon Cancer into Molecular Subtypes: Characterization, Validation, and Prognostic Value. 10: e1001453, 2013.

### Figure S1

##### Clone #13

**CRISPR guide sequence**

135 AGGAGAGTGTGCTGG**GCAGACGATGCTGGACACGA**TGGAGGCGCCCGGCC 184

AGGAGAGTGTGCTGGGCAG------------ACGATGGAGGCGCCCGGCC

AGGAGAGTGTGCTGGGCAGACGATGCTGGACA-GATGGAGGCGCCCGGCC

##### Clone #20

135 AGGAGAGTGTGCTGG**GCAGACGATGCTGGACACGA**TGGAGGCGCCCGGCC 184

AGGAGAGTGTGCTGGGCAGACGATGCTGGAC-CGATGGAGGCGCCCGGCC

AGGAGAGTGTGCTGGGCAGACGATGCTGGACA-GATGGAGGCGCCCGGCC

##### Clone #25

135 AGGAGAGTGTGCTGG**GCAGACGATGCTGGACACGA**TGGAGGCGCCCGGCC 184

AGGAGAGTGTGCTGGGCAGACGATGCTGGAC-CGATGGAGGCGCCCGGCC

AGGAGAGTGTGCTGGGCAGACGATGCTGGACA-GATGGAGGCGCCCGGCC

### Figure S2

#### Dataset: GSE37892

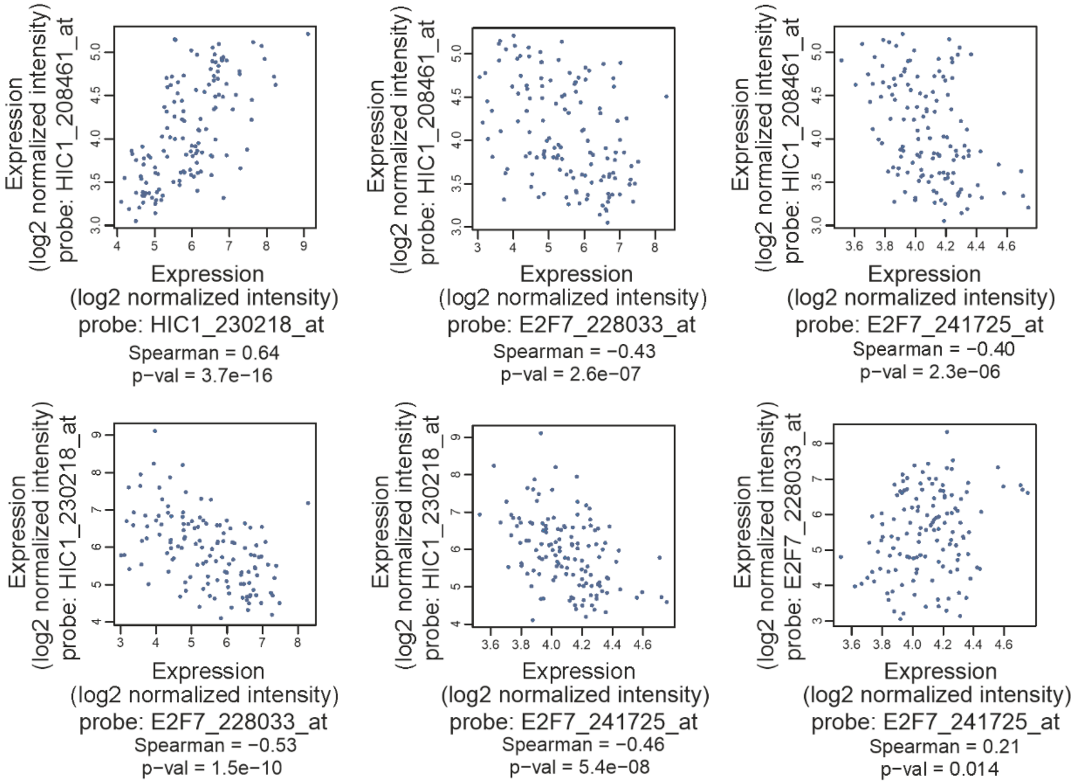


#### Dataset: GSE39582

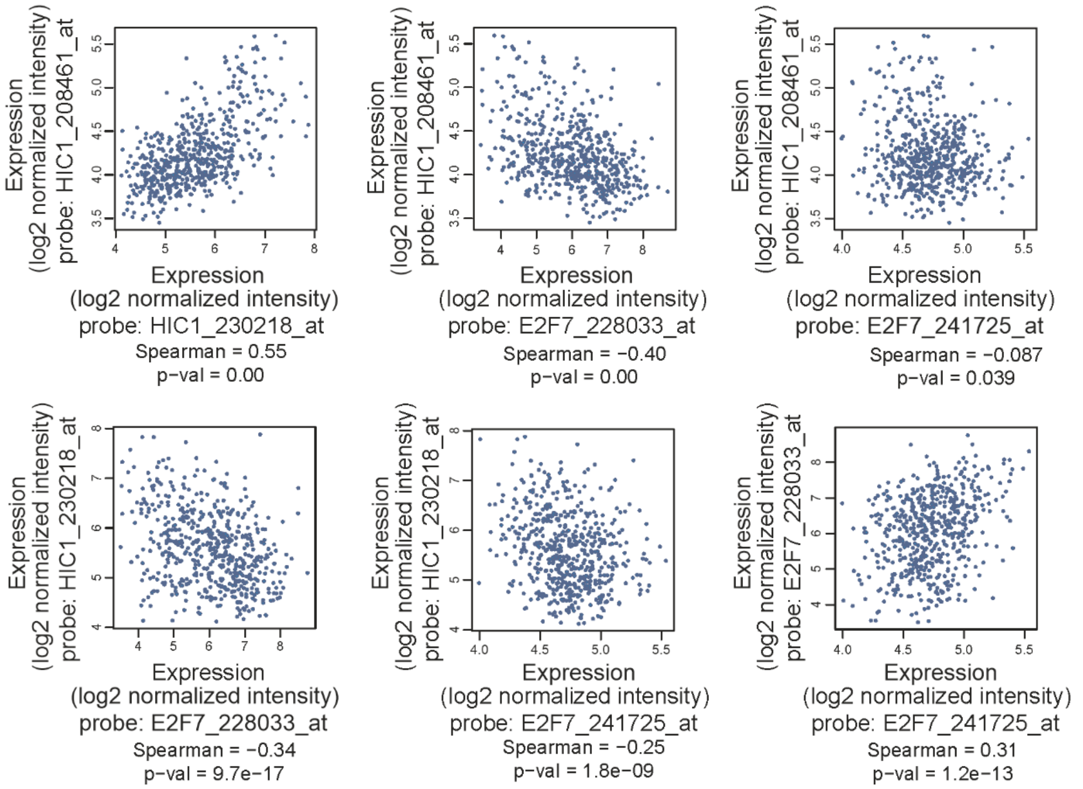

#### Figure S3

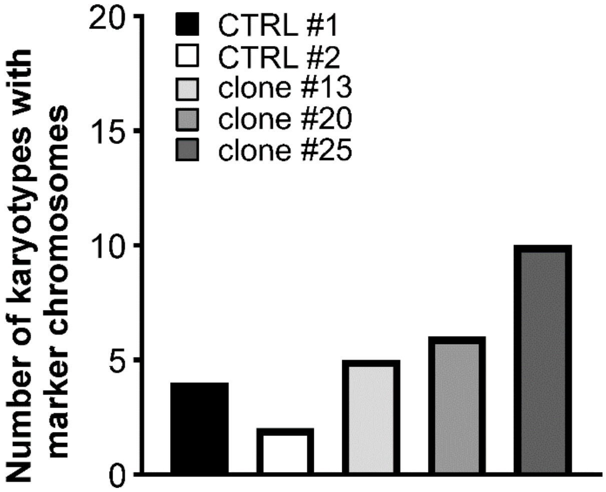

Number of karyotypes with marker chromosomes
20
15
10
5
0
CTRL #1
CTRL #2
clone #13
clone #20
clone #25

#### Tables S1. List of primers and probes used in the study

**qRT-PCR primers**

| Gene symbol | Organism | Sequence (5´ to 3´) | Application | Probe |
|---|---|---|---|---|
| *ACTB* | Human | Forward: GGCATCCTCACCCTGAAGTA Reverse: AGGTGTGGTGCCAGATTTTC | qRT-PCR | |
| *E2F7* | Human | Forward: GGCATCCTCACCCTGAAGTA Reverse: TAGTCTGGGCTGCCTGGGGC | qRT-PCR | |
| *E2F7* | Human | Forward: GGCATCCTCACCCTGAAGTA Reverse: CTAAAGAGTAGCCACCTGATCCTT | UPL-based* qRT-PCR | 6 |
| *HIC1* | Human | Forward: CTGAGACGCGACCAGGAC Reverse: CAGCACACTCTCCTGGGG | qRT-PCR | |
| *HIC1* | Human | Forward: CTGAGACGCGACCAGGAC Reverse: CTTGGTGCGCTGGTTGTT | UPL-based qRT-PCR | 27 |
| *TBP* | Human | Forward: GAACATCATGGATCAGAACAACA Reverse: ATAGGGATTCCGGGAGTCAT | UPL-based qRT-PCR | 87 |
| *UBB* | Human | Forward: GCTTTGTTGGGTGAGCTTGT Reverse: TCACGAAGATCTGCATTTTGA | qRT-PCR | |
| *UBB* | Human | Forward: AGGATCCTGGTATCCGCTAAC Reverse: TCACATTTTCGATGGTGTCACT | UPL-based qRT-PCR | 39 |

***E2F7-luc reporter***

| Primer name | Sequence (5´ to 3´) |
|---|---|
| *FhE2F7* _LL | GGGTGAAATTGGGTGCTGAA |
| *RhE2F7* _LL | TCGTAGTCCCCGCTAAAACA |

***CRISPR/Cas9 editing oligos***

| Primer name | Sequence (5´ to 3´) |
|---|---|
| *FhHIC1* crispr | caccGCAGACGATGCTGGACACGA |
| *RhHIC1* crispr | aaacTCGTGTCCAGCATCGTCTGC |
| *FhHIC1 pARv-RFP* | GCAGACGATGCTGGACACGAtggAT |
| *RhHIC1 pARv-RFP* | ccaTCGTGTCCAGCATCGTCTGC |

***HIC1* genotyping PCR primers**

| Primer name | | Sequence (5´ to 3´) |
|---|---|---|
| *FhHIC1* crispr test | Human | GAAGGGGAAGTGGAGGGGAGAAGTG |
| *FhHIC1* crispr test | Human | GATGATCACGTCGCACAAGAAGCCC |

**Sequencing primers**

| Vector background | Sequence (5´ to 3´) |
|---|---|
| pARv-RFP | CACCGCTAATTCAAAGCAACCG |
| lentiCRISPRv2 | CTACTATTCTTTCCCCTGCACTGTAC |
| pGEM T-easy | GATTTAGGTGACACTATAG |

*) UPL, Universal ProbeLibrary (Roche Life Sciences)

### Table S2. Genes differentially expressed in WI38 cells treated with HIC1 siRNA 48 or 72 hours after transfection

**A combined list of 158 genes differentially expressed in WI38 cells treated with *HIC1* siRNA from Ambion and Dharmacon companies vs. control siRNA 48 hours upon transfection; adjusted p-value < 0.05 and |Log FC| ≥ 1**

| ENTREZ No. | Symbol | Gene name | Log FC | |
|---|---|---|---|---|
| | | | Dhar 48h | Amb 48h |
| 3920 | LAMP2 | lysosomal-associated membrane protein 2 | 2.96 | 2.66 |
| 54499 | TMCO1 | transmembrane and coiled-coil domains 1 | 2.46 | 2.13 |
| 79191 | IRX3 | iroquois homeobox 3 | 2.32 | 1.08 |
| 1009 | CDH11 | cadherin 11, type 2, OB-cadherin (osteoblast) | 2.28 | 2.39 |
| 5649 | RELN | reelin | 2.11 | 2.92 |
| 94241 | TP53INP1 | tumor protein p53 inducible nuclear protein 1 | 2.01 | 1.25 |
| 8942 | KYNU | kynureninase | 1.98 | 2.39 |
| 94081 | SFXN1 | sideroflexin 1 | 1.97 | 2.17 |
| 4507 | MTAP | methylthioadenosine phosphorylase | 1.84 | 1.46 |
| 3915 | LAMC1 | laminin, gamma 1 (formerly LAMB2) | 1.8 | 2.21 |
| 83700 | JAM3 | junctional adhesion molecule 3 | 1.75 | 2.1 |
| 64979 | MRPL36 | mitochondrial ribosomal protein L36 | 1.75 | 1.7 |
| 116028 | RMI2 | RecQ mediated genome instability 2 | 1.75 | 1.19 |
| 11145 | PLA2G16 | phospholipase A2, group XVI | 1.72 | 1.11 |
| 79974 | CPED1 | cadherin-like and PC-esterase domain containing 1 | 1.66 | 2.24 |
| 138151 | NACC2 | NACC family member 2, BEN and BTB (POZ) domain containing | 1.66 | 1.95 |
| 6836 | SURF4 | surfeit 4 | 1.64 | 2.12 |
| 57590 | WDFY1 | WD repeat and FYVE domain containing 1 | 1.63 | 2.28 |
| 55062 | WIPI1 | WD repeat domain, phosphoinositide interacting 1 | 1.61 | 2.23 |
| 88455 | ANKRD13A | ankyrin repeat domain 13A | 1.59 | 1.98 |
| 144455 | E2F7 | E2F transcription factor 7 | 1.59 | 2.28 |
| 10778 | ZNF271 | zinc finger protein 271 | 1.58 | 1.7 |
| 64423 | INF2 | inverted formin, FH2 and WH2 domain containing | 1.55 | 1.14 |
| 51449 | PCYOX1 | prenylcysteine oxidase 1 | 1.55 | 1.32 |
| 5894 | RAF1 | v-raf-1 murine leukemia viral oncogene homolog 1 | 1.55 | 2.06 |
| 55437 | STRADB | STE20-related kinase adaptor beta | 1.55 | 1.91 |
| 29904 | EEF2K | eukaryotic elongation factor-2 kinase | 1.54 | 1.4 |
| 10049 | DNAJB6 | DnaJ (Hsp40) homolog, subfamily B, member 6 | 1.52 | 1.52 |
| 79443 | FYCO1 | FYVE and coiled-coil domain containing 1 | 1.51 | 3.33 |
| 665 | BNIP3L | BCL2/adenovirus E1B 19kDa interacting protein 3-like | 1.5 | 2.36 |
| 134147 | CMBL | carboxymethylenebutenolidase homolog (Pseudomonas) | 1.5 | 1.13 |
| 11041 | B3GNT1 | UDP-GlcNAc:betaGal beta-1,3-N-acetylglucosaminyltransferase 1 | 1.43 | 1.23 |
| 900 | CCNG1 | cyclin G1 | 1.41 | 1.61 |
| 64682 | ANAPC1 | anaphase promoting complex subunit 1 | 1.39 | 1.98 |
| 4343 | MOV10 | Mov10, Moloney leukemia virus 10, homolog (mouse) | 1.39 | 1.08 |
| 79710 | MORC4 | MORC family CW-type zinc finger 4 | 1.37 | 1.58 |
| 23433 | RHOQ | ras homolog family member Q | 1.37 | 1.6 |
| 148266 | ZNF569 | zinc finger protein 569 | 1.34 | 1.17 |
| 57089 | ENTPD7 | ectonucleoside triphosphate diphosphohydrolase 7 | 1.32 | 1.3 |
| 23034 | SAMD4A | sterile alpha motif domain containing 4A | 1.31 | 1.58 |
| 6542 | SLC7A2 | solute carrier family 7 (cationic amino acid transporter, y+ system), member 2 | 1.31 | 2.73 |
| 54902 | TTC19 | tetratricopeptide repeat domain 19 | 1.31 | 1 |
| 4090 | SMAD5 | SMAD family member 5 | 1.3 | 1 |
| 57700 | FAM160B1 | family with sequence similarity 160, member B1 | 1.29 | 1.53 |
| 317 | APAF1 | apoptotic peptidase activating factor 1 | 1.27 | 1.6 |
| 64764 | CREB3L2 | cAMP responsive element binding protein 3-like 2 | 1.27 | 1.77 |

| | | | | |
|---|---|---|---|---|
| 132671 | SPATA18 | spermatogenesis associated 18 | 1.27 | 1.49 |
| 3930 | LBR | lamin B receptor | 1.26 | 1.27 |
| 301 | ANXA1 | annexin A1 | 1.25 | 2.11 |
| 55603 | FAM46A | family with sequence similarity 46, member A | 1.24 | 2.06 |
| 24140 | FTSJ1 | FtsJ RNA methyltransferase homolog 1 (E. coli) | 1.24 | 1.47 |
| 114904 | C1QTNF6 | C1q and tumor necrosis factor related protein 6 | 1.23 | 1.12 |
| 5156 | PDGFRA | platelet-derived growth factor receptor, alpha polypeptide | 1.22 | 1.17 |
| 56935 | SMCO4 | single-pass membrane protein with coiled-coil domains 4 | 1.2 | 1.32 |
| 4976 | OPA1 | optic atrophy 1 (autosomal dominant) | 1.19 | 1.2 |
| 27346 | TMEM97 | transmembrane protein 97 | 1.17 | 1.72 |
| 4216 | MAP3K4 | mitogen-activated protein kinase kinase kinase 4 | 1.15 | 1.59 |
| 124540 | MSI2 | musashi RNA-binding protein 2 | 1.15 | 1.67 |
| 118429 | ANTXR2 | anthrax toxin receptor 2 | 1.14 | 1.88 |
| 5010 | CLDN11 | claudin 11 | 1.14 | 1.46 |
| 90317 | ZNF616 | zinc finger protein 616 | 1.12 | 1.04 |
| 80777 | CYB5B | cytochrome b5 type B (outer mitochondrial membrane) | 1.11 | 1.01 |
| 9459 | ARHGEF6 | Rac/Cdc42 guanine nucleotide exchange factor (GEF) 6 | 1.1 | 1.76 |
| 1514 | CTSL | cathepsin L | 1.1 | 1.4 |
| 253558 | LCLAT1 | lysocardiolipin acyltransferase 1 | 1.09 | 1.11 |
| 8434 | RECK | reversion-inducing-cysteine-rich protein with kazal motifs | 1.09 | 1.62 |
| 60560 | NAA35 | N(alpha)-acetyltransferase 35, NatC auxiliary subunit | 1.08 | 1.11 |
| 84236 | RHBDD1 | rhomboid domain containing 1 | 1.08 | 1.85 |
| 79646 | PANK3 | pantothenate kinase 3 | 1.07 | 1.06 |
| 56666 | PANX2 | pannexin 2 | 1.07 | 1.15 |
| 10974 | ADIRF | adipogenesis regulatory factor | 1.05 | 1.09 |
| 221336 | BEND6 | BEN domain containing 6 | 1.04 | 1.42 |
| 5654 | HTRA1 | HtrA serine peptidase 1 | 1.04 | 1.14 |
| 10443 | N4BP2L2 | NEDD4 binding protein 2-like 2 | 1.04 | 1.18 |
| 7008 | TEF | thyrotrophic embryonic factor | 1.04 | 1.1 |
| 54918 | CMTM6 | CKLF-like MARVEL transmembrane domain containing 6 | 1.03 | 1.18 |
| 1290 | COL5A2 | collagen, type V, alpha 2 | 1.03 | 1.5 |
| 25940 | FAM98A | family with sequence similarity 98, member A | 1.03 | 1.69 |
| 6745 | SSR1 | signal sequence receptor, alpha | 1.02 | 1.31 |
| 79986 | ZNF702P | zinc finger protein 702, pseudogene | 1 | 1.53 |
| 4204 | MECP2 | methyl CpG binding protein 2 (Rett syndrome) | -1 | -1.96 |
| 861 | RUNX1 | runt-related transcription factor 1 | -1 | -1.26 |
| 65997 | RASL11B | RAS-like, family 11, member B | -1.01 | -1.07 |
| 4155 | MBP | myelin basic protein | -1.02 | -1.48 |
| 79144 | PPDPF | pancreatic progenitor cell differentiation and proliferation factor | -1.04 | -1.04 |
| 51642 | MRPL48 | mitochondrial ribosomal protein L48 | -1.05 | -1.01 |
| 6347 | CCL2 | chemokine (C-C motif) ligand 2 | -1.06 | -1.48 |
| 3430 | IFI35 | interferon-induced protein 35 | -1.07 | -1.26 |
| 56998 | CTNNBIP1 | catenin, beta interacting protein 1 | -1.08 | -1.73 |
| 338773 | TMEM119 | transmembrane protein 119 | -1.09 | -2.39 |
| 3212 | HOXB2 | homeobox B2 | -1.1 | -1.74 |
| 9519 | TBPL1 | TBP-like 1 | -1.1 | -1.92 |
| 2634 | GBP2 | guanylate binding protein 2, interferon-inducible | -1.12 | -1.68 |
| 64965 | MRPS9 | mitochondrial ribosomal protein S9 | -1.12 | -1.01 |
| 57403 | RAB22A | RAB22A, member RAS oncogene family | -1.12 | -1.14 |
| 10437 | IFI30 | interferon, gamma-inducible protein 30 | -1.13 | -1.57 |
| 857 | CAV1 | caveolin 1, caveolae protein, 22kDa | -1.19 | -1.02 |
| 3669 | ISG20 | interferon stimulated exonuclease gene 20kDa | -1.19 | -1.21 |
| 10653 | SPINT2 | serine peptidase inhibitor, Kunitz type, 2 | -1.21 | -1.08 |

| | | | | |
|---|---|---|---|---|
| 51606 | ATP6V1H | ATPase, H+ transporting, lysosomal 50/57kDa, V1 subunit H | -1.24 | -1.14 |
| 3728 | JUP | junction plakoglobin | -1.24 | -1.25 |
| 58499 | ZNF462 | zinc finger protein 462 | -1.25 | -1.25 |
| 2878 | GPX3 | glutathione peroxidase 3 (plasma) | -1.27 | -2.1 |
| 11046 | SLC35D2 | solute carrier family 35 (UDP-GlcNAc/UDP-glucose transporter), member D2 | -1.27 | -1.19 |
| 115572 | FAM46B | family with sequence similarity 46, member B | -1.3 | -1.01 |
| 3006 | HIST1H1C | histone cluster 1, H1c | -1.33 | -1.02 |
| 8566 | PDXK | pyridoxal (pyridoxine, vitamin B6) kinase | -1.33 | -1.42 |
| 2787 | GNG5 | guanine nucleotide binding protein (G protein), gamma 5 | -1.34 | -2.02 |
| 6665 | SOX15 | SRY (sex determining region Y)-box 15 | -1.34 | -1.51 |
| 120 | ADD3 | adducin 3 (gamma) | -1.35 | -1.49 |
| 1839 | HBEGF | heparin-binding EGF-like growth factor | -1.35 | -1.18 |
| 5155 | PDGFB | platelet-derived growth factor beta polypeptide | -1.35 | -1.37 |
| 624 | BDKRB2 | bradykinin receptor B2 | -1.39 | -1.2 |
| 3148 | HMGB2 | high mobility group box 2 | -1.39 | -1.35 |
| 5292 | PIM1 | pim-1 oncogene | -1.4 | -1.38 |
| 94240 | EPSTI1 | epithelial stromal interaction 1 (breast) | -1.42 | -1.52 |
| 11098 | PRSS23 | protease, serine, 23 | -1.42 | -1.56 |
| 55905 | RNF114 | ring finger protein 114 | -1.44 | -1.07 |
| 797 | CALCB | calcitonin-related polypeptide beta | -1.45 | -1.41 |
| 23531 | MMD | monocyte to macrophage differentiation-associated | -1.47 | -1.04 |
| 169611 | OLFML2A | olfactomedin-like 2A | -1.48 | -1.4 |
| 29890 | RBM15B | RNA binding motif protein 15B | -1.48 | -1.49 |
| 7336 | UBE2V2 | ubiquitin-conjugating enzyme E2 variant 2 | -1.48 | -1.73 |
| 3107 | HLA-C | major histocompatibility complex, class I, C | -1.5 | -1.01 |
| 26301 | GBGT1 | globoside alpha-1,3-N-acetylgalactosaminyltransferase 1 | -1.51 | -1.73 |
| 132946 | ARL9 | ADP-ribosylation factor-like 9 | -1.55 | -1.89 |
| 836 | CASP3 | caspase 3, apoptosis-related cysteine peptidase | -1.56 | -1.27 |
| 2181 | ACSL3 | acyl-CoA synthetase long-chain family member 3 | -1.61 | -1.06 |
| 652 | BMP4 | bone morphogenetic protein 4 | -1.62 | -2.6 |
| 306 | ANXA3 | annexin A3 | -1.64 | -1.19 |
| 9249 | DHRS3 | dehydrogenase/reductase (SDR family) member 3 | -1.66 | -1.42 |
| 3589 | IL11 | interleukin 11 | -1.7 | -1.8 |
| 5217 | PFN2 | profilin 2 | -1.71 | -1.44 |
| 6447 | SCG5 | secretogranin V (7B2 protein) | -1.72 | -1.45 |
| 27075 | TSPAN13 | tetraspanin 13 | -1.74 | -1.07 |
| 89796 | NAV1 | neuron navigator 1 | -1.76 | -1.18 |
| 11178 | LZTS1 | leucine zipper, putative tumor suppressor 1 | -1.81 | -1.64 |
| 11332 | ACOT7 | acyl-CoA thioesterase 7 | -1.85 | -1.81 |
| 152007 | GLIPR2 | GLI pathogenesis-related 2 | -1.92 | -1.5 |
| 91283 | MSANTD3 | Myb/SANT-like DNA-binding domain containing 3 | -1.93 | -1.42 |
| 3976 | LIF | leukemia inhibitory factor | -1.99 | -1.77 |
| 6236 | RRAD | Ras-related associated with diabetes | -2.01 | -1.22 |
| 1134 | CHRNA1 | cholinergic receptor, nicotinic, alpha 1 (muscle) | -2.02 | -1.35 |
| 3875 | KRT18 | keratin 18 | -2.11 | -1.27 |
| 23327 | NEDD4L | neural precursor cell expressed, developmentally down-regulated 4-like, E3 ubiquitin protein ligase | -2.13 | -2.48 |
| 9289 | GPR56 | G protein-coupled receptor 56 | -2.18 | -1.17 |
| 94234 | FOXQ1 | forkhead box Q1 | -2.22 | -1.01 |
| 4908 | NTF3 | neurotrophin 3 | -2.23 | -1.29 |
| 4088 | SMAD3 | SMAD family member 3 | -2.35 | -1.64 |
| 7292 | TNFSF4 | tumor necrosis factor (ligand) superfamily, member 4 | -2.35 | -1.24 |
| 6385 | SDC4 | syndecan 4 | -2.36 | -2.61 |
| 56180 | MOSPD1 | motile sperm domain containing 1 | -2.4 | -1.28 |

| | | | | |
|---|---|---|---|---|
| 221091 | LRRN4CL | LRRN4 C-terminal like | -2.46 | -1.63 |
| 5743 | PTGS2 | prostaglandin-endoperoxide synthase 2 (prostaglandin G/H synthase and cyclooxygenase) | -2.65 | -1.47 |
| 8869 | ST3GAL5 | ST3 beta-galactoside alpha-2,3-sialyltransferase 5 | -2.71 | -1.37 |
| 3569 | IL6 | interleukin 6 (interferon, beta 2) | -2.82 | -1.8 |
| 1734 | DIO2 | deiodinase, iodothyronine, type II | -3.01 | -1.18 |
| 3045 | HBD | hemoglobin, delta | -3.53 | -3.14 |

#### Table S3. Genes differentially expressed in WI38 cells treated with HIC1 siRNA at 48- and 72-hour time intervals

| ENTREZ No. | Symbol | Gene name | Log FC | | | |
|---|---|---|---|---|---|---|
| | | | Dhar 48h | Amb 48h | Dhar 72h | Amb 72h |
| 2181 | ACSL3 | acyl-CoA synthetase long-chain family member 3 | -1.61 | -1.06 | -1.47 | -1.24 |
| 120 | ADD3 | adducin 3 (gamma) | -1.35 | -1.49 | -1.06 | -1.15 |
| 10974 | ADIRF | adipogenesis regulatory factor | 1.05 | 1.09 | 1.11 | 1.9 |
| 132946 | ARL9 | ADP-ribosylation factor-like 9 | -1.55 | -1.89 | -1.28 | -1.12 |
| 301 | ANXA1 | annexin A1 | 1.25 | 2.11 | 1.21 | 2.03 |
| 306 | ANXA3 | annexin A3 | -1.64 | -1.19 | -1.84 | -1.12 |
| 665 | BNIP3L | BCL2/adenovirus E1B 19kDa interacting protein 3-like | 1.5 | 2.36 | 1.71 | 1.66 |
| 652 | BMP4 | bone morphogenetic protein 4 | -1.62 | -2.6 | -1.68 | -3.18 |
| 79974 | CPED1 | cadherin-like and PC-esterase domain containing 1 | 1.66 | 2.24 | 1.48 | 1.61 |
| 56998 | CTNNBIP1 | catenin, beta interacting protein 1 | -1.08 | -1.73 | -1.06 | -1.71 |
| 1134 | CHRNA1 | cholinergic receptor, nicotinic, alpha 1 (muscle) | -2.02 | -1.35 | -1.86 | -1.45 |
| 900 | CCNG1 | cyclin G1 | 1.41 | 1.61 | 1.39 | 1.15 |
| 9249 | DHRS3 | dehydrogenase/reductase (SDR family) member 3 | -1.66 | -1.42 | -1.72 | -1.3 |
| 144455 | E2F7 | E2F transcription factor 7 | 1.59 | 2.28 | 1.96 | 1.53 |
| 115572 | FAM46B | family with sequence similarity 46, member B | -1.3 | -1.01 | -2.24 | -1.64 |
| 79443 | FYCO1 | FYVE and coiled-coil domain containing 1 | 1.51 | 3.33 | 1.76 | 2.52 |
| 9289 | GPR56 | G protein-coupled receptor 56 | -2.18 | -1.17 | -2.34 | -1.03 |
| 152007 | GLIPR2 | GLI pathogenesis-related 2 | -1.92 | -1.5 | -2.04 | -1.8 |
| 26301 | GBGT1 | globoside alpha-1,3-N-acetylgalactosaminyltransferase 1 | -1.51 | -1.73 | -1.21 | -1.34 |
| 2878 | GPX3 | glutathione peroxidase 3 (plasma) | -1.27 | -2.1 | -1.32 | -1.61 |
| 2787 | GNG5 | guanine nucleotide binding protein (G protein), gamma 5 | -1.34 | -2.02 | -1.04 | -1.08 |
| 2634 | GBP2 | guanylate binding protein 2, interferon-inducible | -1.12 | -1.68 | -1.44 | -2.43 |
| 3045 | HBD | hemoglobin, delta | -3.53 | -3.14 | -4.82 | -3.76 |
| 3148 | HMGB2 | high mobility group box 2 | -1.39 | -1.35 | -1.89 | -1.3 |
| 5654 | HTRA1 | HtrA serine peptidase 1 | 1.04 | 1.14 | 1.05 | 1.03 |
| 10437 | IFI30 | interferon, gamma-inducible protein 30 | -1.13 | -1.57 | -1.6 | -1.33 |
| 3430 | IFI35 | interferon-induced protein 35 | -1.07 | -1.26 | -1.11 | -1.09 |
| 3589 | IL11 | interleukin 11 | -1.7 | -1.8 | -1.12 | -1.37 |
| 3569 | IL6 | interleukin 6 (interferon, beta 2) | -2.82 | -1.8 | -1.94 | -1.86 |
| 3875 | KIAA0930 | KIAA0930 | -1.63 | -1.7 | -1.43 | -1.46 |
| 3875 | KRT18 | keratin 18 | -2.11 | -1.27 | -1.51 | -1.33 |
| 8942 | KYNU | kynureninase | 1.98 | 2.39 | 2.2 | 2.92 |
| 3915 | LAMC1 | laminin, gamma 1 (formerly LAMB2) | 1.8 | 2.21 | 1.54 | 1.08 |
| 11178 | LZTS1 | leucine zipper, putative tumor suppressor 1 | -1.81 | -1.64 | -2.32 | -1.99 |
| 3976 | LIF | leukemia inhibitory factor | -1.99 | -1.77 | -1.82 | -1.64 |
| 221091 | LRRN4CL | LRRN4 C-terminal like | -2.46 | -1.63 | -2.76 | -1.69 |
| 3920 | LAMP2 | lysosomal-associated membrane protein 2 | 2.96 | 2.66 | 2.27 | 1.53 |
| 64979 | MRPL36 | mitochondrial ribosomal protein L36 | 1.75 | 1.7 | 1.42 | 1.7 |
| 23531 | MMD | monocyte to macrophage differentiation-associated | -1.47 | -1.04 | -1.39 | -1.01 |
| 138151 | NACC2 | NACC family member 2, BEN and BTB (POZ) domain containing | 1.66 | 1.95 | 1.42 | 1.98 |
| 23327 | NEDD4L | neural precursor cell expressed, developmentally down-regulated 4-like, E3 ubiquitin protein ligase | -2.13 | -2.48 | -1.48 | -1.62 |
| 89796 | NAV1 | neuron navigator 1 | -1.76 | -1.18 | -1.07 | -1.13 |
| 169611 | OLFML2A | olfactomedin-like 2A | -1.48 | -1.4 | -2.34 | -1.93 |
| 56666 | PANX2 | pannexin 2 | 1.07 | 1.15 | 1.15 | 1.13 |
| 11145 | PLA2G16 | phospholipase A2, group XVI | 1.72 | 1.11 | 1.72 | 1.68 |
| 5292 | PIM1 | pim-1 oncogene | -1.4 | -1.38 | -2.17 | -2 |

| | | | | | | |
|---|---|---|---|---|---|---|
| 5155 | PDGFB | platelet-derived growth factor beta polypeptide | -1.35 | -1.37 | -1.26 | -1.35 |
| 11098 | PRSS23 | protease, serine, 23 | -1.42 | -1.56 | -1.43 | -2.27 |
| 8566 | PDXK | pyridoxal (pyridoxine, vitamin B6) kinase | -1.33 | -1.42 | -1.5 | -1.49 |
| 23433 | RHOQ | ras homolog family member Q | 1.37 | 1.6 | 1.09 | 1.34 |
| 5649 | RELN | reelin | 2.11 | 2.92 | 1.18 | 1.63 |
| 390 | RND3 | Rho family GTPase 3 | -1.7 | 1.85 | -1.07 | 1.37 |
| 84236 | RHBDD1 | rhomboid domain containing 1 | 1.08 | 1.85 | 1.01 | 1.19 |
| 29890 | RBM15B | RNA binding motif protein 15B | -1.48 | -1.49 | -1.2 | -1.1 |
| 10653 | SPINT2 | serine peptidase inhibitor, Kunitz type, 2 | -1.21 | -1.08 | -1.14 | -1.35 |
| 94081 | SFXN1 | sideroflexin 1 | 1.97 | 2.17 | 1.28 | 1.25 |
| 4088 | SMAD3 | SMAD family member 3 | -2.35 | -1.64 | -1.85 | -1.52 |
| 132671 | SPATA18 | spermatogenesis associated 18 | 1.27 | 1.49 | 2.01 | 1.08 |
| 6665 | SOX15 | SRY (sex determining region Y)-box 15 | -1.34 | -1.51 | -1.38 | -1.59 |
| 8869 | ST3GAL5 | ST3 beta-galactoside alpha-2,3-sialyltransferase 5 | -2.71 | -1.37 | -1.35 | -1.14 |
| 23034 | SAMD4A | sterile alpha motif domain containing 4A | 1.31 | 1.58 | 1.67 | 1.29 |
| 6836 | SURF4 | surfeit 4 | 1.64 | 2.12 | 1.03 | 1.15 |
| 6385 | SDC4 | syndecan 4 | -2.36 | -2.61 | -1.74 | -1.86 |
| 54499 | TMCO1 | transmembrane and coiled-coil domains 1 | 2.46 | 2.13 | 2.2 | 2.12 |
| 338773 | TMEM119 | transmembrane protein 119 | -1.09 | -2.39 | -1.23 | -2.15 |
| 94241 | TP53INP1 | tumor protein p53 inducible nuclear protein 1 | 2.01 | 1.25 | 2.35 | 1.34 |
| 7336 | UBE2V2 | ubiquitin-conjugating enzyme E2 variant 2 | -1.48 | -1.73 | -1.1 | -1.12 |
| 5894 | RAF1 | v-raf-1 murine leukemia viral oncogene homolog 1 | 1.55 | 2.06 | 1.19 | 1.18 |
| 57590 | WDFY1 | WD repeat and FYVE domain containing 1 | 1.63 | 2.28 | 1.41 | 1.78 |
| 55062 | WIPI1 | WD repeat domain, phosphoinositide interacting 1 | 1.61 | 2.23 | 1.56 | 2.24 |
| 10778 | ZNF271 | zinc finger protein 271 | 1.58 | 1.7 | 1.18 | 1.12 |
| 79986 | ZNF702P | zinc finger protein 702, pseudogene | 1 | 1.53 | 1.3 | 1.14 |
